# A systematic review and meta-analysis of tau phosphorylation in mouse models of familial Alzheimer’s disease

**DOI:** 10.1101/2023.10.16.562481

**Authors:** Malamati Kourti, Athanasios Metaxas

## Abstract

**Background:** Transgenic models of familial Alzheimer’s disease (AD) serve as valuable tools for probing the molecular mechanisms associated with amyloid-beta (Aβ)-induced pathology. Here, we sought to evaluate the levels of phosphorylated tau (p-tau) protein, and explore potential age-related variations in the hyperphosphorylation of tau, in mouse models of cerebral amyloidosis.

**Methods:** The PubMed and Scopus databases were searched for studies measuring soluble p-tau in 5xFAD, *APP*_swe_/*PSEN1*_de9_, J20 and APP23 mice. Data were extracted and analyzed using standardized procedures.

**Results:** For the 5xFAD model, the search yielded 36 studies eligible for meta-analysis. Levels of p-tau were higher in 5xFAD mice relative to control, a difference that was evident in both the carboxy-terminal (CT) and proline-rich (PR) domains of tau. Age negatively moderated the effects of genotype on CT domain phosphorylated tau, particularly in studies using hybrid mice, female mice, and preparations from the cortex. For the *APP*_swe_/*PSEN1*_de9_ model, the search yielded 27 studies. Analysis showed tau hyperphosphorylation in transgenic vs. control animals, evident in both the CT and PR regions of tau. Age positively moderated the effects of genotype on PR domain phosphorylated tau in the cortex of *APP*_swe_/*PSEN1*_de9_ mice. A meta-analysis was not performed for the J20 and APP23 models, due to the limited number of studies measuring p-tau levels in these mice (<10 studies).

**Conclusions:** Although tau is hyperphosphorylated in both 5xFAD and *APP*_swe_/*PSEN1*_de9_ mice, the effects of ageing on p-tau are contingent upon the mouse model being examined. These observations emphasize the importance of tailoring model selection to the appropriate disease stage when assessing the relationship between Aβ and tau, and suggest that there are optimal intervention points for the administration of both anti-amyloid and anti-tau therapies.

## 1. Introduction

Alzheimer’s disease (AD) is the most common cause of dementia, accounting for approximately 60-80% of all dementia cases in people over the age of 65 (Alzheimer’s Association, 2023). AD is a chronic, irreversible, degenerative disorder that is characterized by the gradual buildup of abnormal protein aggregates in the brain, most notably plaques and tangles. Plaques accumulate extraneuronally and are primarily composed of amyloid-β (Aβ) peptides, which are formed by the proteolytic cleavage of the amyloid precursor protein (APP). Neurofibrillary tangles (NFTs) accumulate within neurons and are primarily composed of hyperphosphorylated MAPT (tau) protein. To date, biomarker evidence for the presence of both Aβ and abnormally phosphorylated (p-tau) soluble and/or fibrillar tau is required for a definitive diagnosis of AD (Jack et al., 2018a).

Recent cross-sectional and longitudinal imaging studies report that neocortical Aβ amyloidosis precedes the accumulation of NFT pathology (Jack et al., 2018b; Jacobs et al., 2018; Lee et al., 2022; Scholl et al., 2016), potentially favouring its propagation from limbic areas to the cerebral cortex in living AD patients. This rather consistent spatiotemporal pattern of propagation places fibrillar Aβ upstream of neocortical tau, and points towards the existence of complex and dynamic links between the two pathognomonic features of AD. In humans, the interplay between Aβ and tau may be disease-stage specific, with increased soluble markers of p-tau181 and p-tau217 exhibiting close correlations with Aβ pathology in the early rather than the later stages of AD (Barthelemy et al., 2020; Mattsson-Carlgren et al., 2020; Ossenkoppele et al., 2021). In transgenic models of familial AD, where Αβ has been shown to enhance tau phenotypes throughout the disease course (Busche and Hyman, 2020), there is evidence to suggest that the functional consequences of the Aβ-tau interaction may also depend on disease stage. Thus, Aβ-induced neuronal excitation is likely to predominate in the early stages, and tau-induced neuronal silencing in the later stages of the disease process (Busche et al., 2019). In both animal models (Gotz et al., 2001; Rasool et al., 2013) and AD patients (Silvestro et al., 2022; van Dyck et al., 2023), removing Aβ by immunotherapy results in decreased tau pathology markers. However, the effectiveness of any anti-Αβ intervention is likely to be higher when given at the early phases of AD (Sims et al., 2023; Sperling et al., 2011). Collectively, these data demonstrate that Aβ and tau interact differentially during the various stages of AD progression, with the early interplay between the soluble forms of Aβ and tau potentially contributing to the pathogenesis of the disorder.

Mouse models of amyloidosis have been engineered to (over)express human/humanized *APP* transgene constructs, carrying single or multiple familial AD mutations, either alone or in combination with mutant human presenilin *(PSEN)* (Sasaguri et al., 2017). To date, more than 100 *APP* and *APP/PSEN* models have been generated, each with unique technical and neuropathological characteristics (Sasaguri et al., 2022). Although most amyloidosis models successfully replicate important AD phenotypes, including progressive cognitive and non-cognitive impairment, age-dependent production and deposition of Aβ, and Aβ-induced microgliosis, astrogliosis and synapse loss, tau pathology has been more difficult to reproduce. In the relatively few studies examining the interaction of Aβ and tau in amyloidosis models, substantial heterogeneity exists in both the extent and the morphological features of pathological tau, across and even within individual models. In *APP*_swe_/*PSEN1*_de9_ mice, for example, plaque-associated neuritic tau has been reported to be devoid of post-translational modifications (Chen et al., 2021), hyperphosphorylated but unable to convert to tangle-like aggregates (Li et al., 2016), as well as in the form of Gallyas-positive, sarkosyl-insoluble filaments (Metaxas et al., 2019).

Considering the dynamic relationship between Aβ and tau, a *prima facie* case for the discrepant patterns of tau pathology in models of familial AD is the extensive variation in the age and disease stage of the animals employed across the preclinical research field. We thus conducted this systematic review and meta-analysis to examine the effects of familial AD mutations on the magnitude of tau phosphorylation in mouse models of amyloidosis, using age as a moderator of the association between genotype and tau pathology. The goal was to obtain a comprehensive picture of the patterns of tau phosphorylation in transgenic mice, enhancing our understanding of the interactions between Aβ and tau at distinct disease stages and informing the ongoing efforts of developing timely therapeutic interventions for AD.

## 2. Materials and Methods

### 2.1 Search strategy and selection criteria

The study has been registered in the international prospective register of systematic reviews (PROSPERO ID: CRD42022342296), and prepared following the methodological recommendations of the Systematic Review Centre for Laboratory Animal Experimentation (SYRCLE) (de Vries et al., 2015). PubMed and Scopus were searched for original research articles up until June 2023. The search strategy consisted of an initial query using the terms *Alzheimer* AND *mouse model* (e.g., *APP*_swe_/*PSEN1*_de9_, 5xFAD etc), followed by a secondary interrogation of the results, using the term *tau* as limit. No publication date restrictions were imposed.

Inclusion criteria were specified as follows: (i) papers were peer-reviewed primary research articles (not reviews) (ii) only mouse models of familial AD were examined, with no restriction on genetic background and sex (no other species included, e.g., transgenic rats) (iii) transgenic and control mice were matched for age (iv) p-tau levels in the brains of transgenic and control mice were determined under baseline conditions by western blot or ELISA. Reasons for exclusion of studies were as follows: (i) articles not in the English language (ii) vital study information missing (e.g., age of experimental animals, n numbers) (iii) qualitative studies (no quantification of p-tau reported) (iv) p-tau measured with methods other than western blot or ELISA (‘not a western blot study’). Additional studies were searched within the reference lists of the selected articles. The eligibility criteria were pilot tested by two reviewers, who independently evaluated the full-length text of the retrieved articles pertaining to the *APP*_swe_/*PSEN1*_dE9_ model. Any differences in opinion regarding article inclusion were resolved by consensus. Subsequently, each reviewer single screened the full-length text of the retrieved articles for a given mouse model (MK: APP23; AM: 5xFAD & J20).

### 2.2 Description of mouse models

A brief description of the models examined, along with the exact search terms used to retrieve information are presented below.

*APP*_swe_/*PSEN1*_dE9_ (OR *APP*_swe_/*PSEN1*_ΔE9_, *APP*_swe_/*PS1*_dE9_, *APP*_swe_/*PS1*_ΔE9_, line 85): These double-transgenic mice carry a chimeric mouse/human *APP* (695) with the Swedish mutation (K670N/M671L) and a mutant human *PSEN1* with the exon-9 deletion variant (dE9), both transgenes directed to central nervous system neurons under the control of the mouse prion protein promoter. Cerebral Aβ plaques are deposited in an age-dependent manner, becoming evident by 6 months of age (Jankowsky et al., 2004; Jankowsky et al., 2001). The mice are available in both hybrid and congenic backgrounds.

5xFAD (OR Tg6799): These mice overexpress five AD-linked mutations, carrying human *APP* (695) transgenes with the Swedish (K670N/M671L), Florida (I716V) and London (V717I) mutations, along with the M146L and L286V mutations in human *PSEN1*. The expression of both transgenes is regulated by the mouse thymocyte differentiation antigen 1 (Thy-1) promoter. Cerebral amyloidosis begins as early as 2 months of age (Oakley et al., 2006). The mice are available in both hybrid and congenic backgrounds.

J20 (OR hAPPJ20, hAβPPSwInd, PDAPP-J20): These mice overexpress human *APP* (695) with the Swedish (K670N/M671L) and the Indiana (V717F) mutations under the control of the human platelet-derived growth factor beta polypeptide (PDGFB). Robust cerebral amyloidosis develops by 5 to 7 months of age (Mucke et al., 2000). The mice are available in the C57BL/6 background.

APP23: These mice overexpress human *APP* (751) with the Swedish (K670N/M671L) mutation under the control of the Thy-1 promoter. Plaques are first observed by 6 months of age (Sturchler-Pierrat et al., 1997). The mice are available in the C57BL/6 background.

### 2.3 Quality assessment

The SYRCLE risk-of-bias (RoB) tool was used to assess the methodological quality of the studies included in this review (Hooijmans et al., 2014). The tool consists of 10 questions, which aim to evaluate the risk of selection, performance, detection, attrition and reporting bias, as well as other biases associated with animal research. All questionnaire entries are phrased in a manner so that a positive answer (“yes”) indicates low RoB, a negative answer (“no”) high RoB, and lack of information (“unclear”) indicates unknown RoB. Three additional items were included to assess reporting bias, namely reporting of any measure of randomization, reporting of any measure of blinding, and reporting of power calculation (Velzen et al., 2021). For these items, a positive answer means reported, and a negative answer not reported. Quality assessment was performed by two independent reviewers for the *APP*_swe_/*PSEN1*_dE9_ mouse model, with differences in opinion resolved by consensus. Individual quality assessments were conducted for APP23, 5xFAD and J20 mice (MK: APP23; AM: 5xFAD & J20).

### 2.4 Data extraction

The information extracted from the selected studies included: (i) study characteristics (number of animals/group, method used for determining tau phosphorylation, phosphorylated epitopes studied, method of euthanasia & brain regions examined) and (ii) animal model characteristics (age at death, sex & background strain).

Tau phosphorylation in AD model mice and age-matched controls was the primary outcome variable extracted for meta-analysis. Where multiple phosphorylation epitopes were reported for an individual sample, mean values were calculated for the entire sample and used for meta-analysis. Data that were reported in graphs were retrieved by using a digital ruler (WebPlotDigitizer, version 4.5, Pacifica, California, USA). Where applicable, authors were contacted for unreported data by email. Standard errors of the mean were transformed into standard deviations (SD). In two studies, missing SD values were imputed based on averaged data within the same study. When the animal group size was reported as a range, the lowest number of animals was used in the analysis. Articles that reported time-dependent changes in tau phosphorylation were included in the review as independent comparisons if different, age-matched groups were used at each time point.

### 2.5 Meta analysis

Exploratory meta-analysis was performed to evaluate the hyperphosphorylation of soluble tau in different mouse models of familial AD. The analysis was performed using the *Meta-Essentials* workbooks for meta-analysis (Suurmond et al., 2017). The standardized mean difference was used as a summary statistic. Hedge’s g and 95% Confidence Intervals (CI) were calculated for each study and presented in a forest plot. Prediction Intervals (PI) were also calculated. Pre-specified analysis included the comparison of tau phosphorylation between transgenic and control mice at the proline-rich (PR; residues 153-243) and the C-terminal domains of tau (CT; residues 368-441; numbering according to the longest TAU isoform), using brain region as a subgroup (cortex vs. hippocampus). *Post-hoc* defined subgroup analysis was performed for sex and background. For all studies, age was used as a predictor of the effect size (moderator). In all cases, a random effects model was used to combine results. Heterogeneity between studies was evaluated by visual inspection of the forest plot and by calculating I^2^. Publication bias was evaluated by constructing funnel plots and performing Egger regression to assess plot asymmetry. The trim & fill method was used to compute bias-adjusted, combined effect sizes. In all cases, statistical significance was set at a<0.05.

## 3. Results

### 3.1 Study selection and characteristics: 5xFAD model

The flowchart of study selection for the 5xFAD model is presented in Fig. 1. Searching in PubMed and Scopus retrieved 970 articles, of which 175 were duplicated between and within databases. After screening the remaining 795 records, a total of 110 primary research articles were assessed for eligibility. Of them, 32 articles did not measure tau phosphorylation compared to control mice, 19 articles reported tau phosphorylation exclusively in a qualitative manner (no quantification), 18 articles quantified tau using methods other than immunoblotting or ELISA, and 5 articles lacked vital study information (e.g., age of animals). The characteristics of the 36 articles that were included in the study are shown in Table 1.

**Fig. 1.**
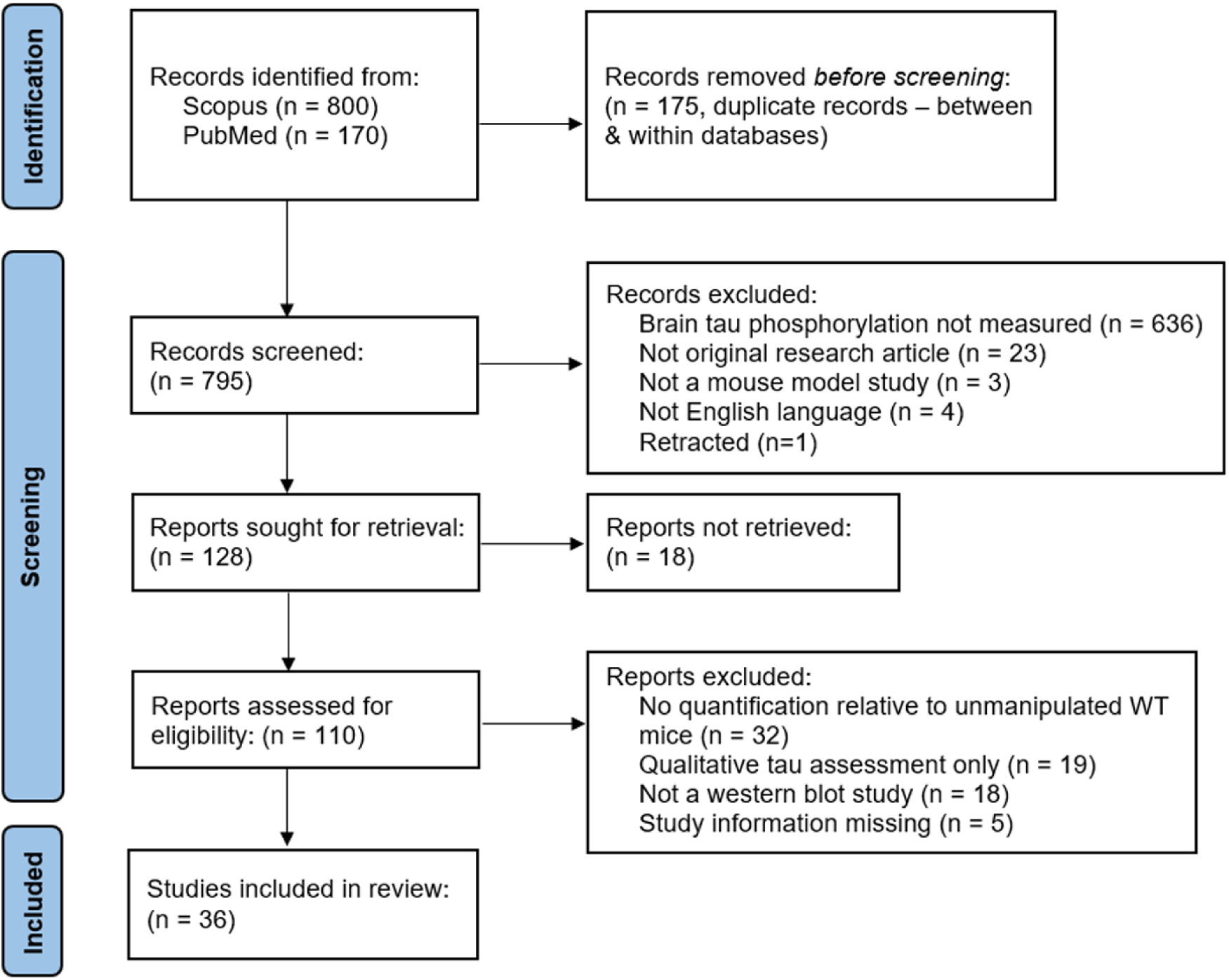
Flowchart of study search and selection for the 5xFAD model.

**Table 1.**
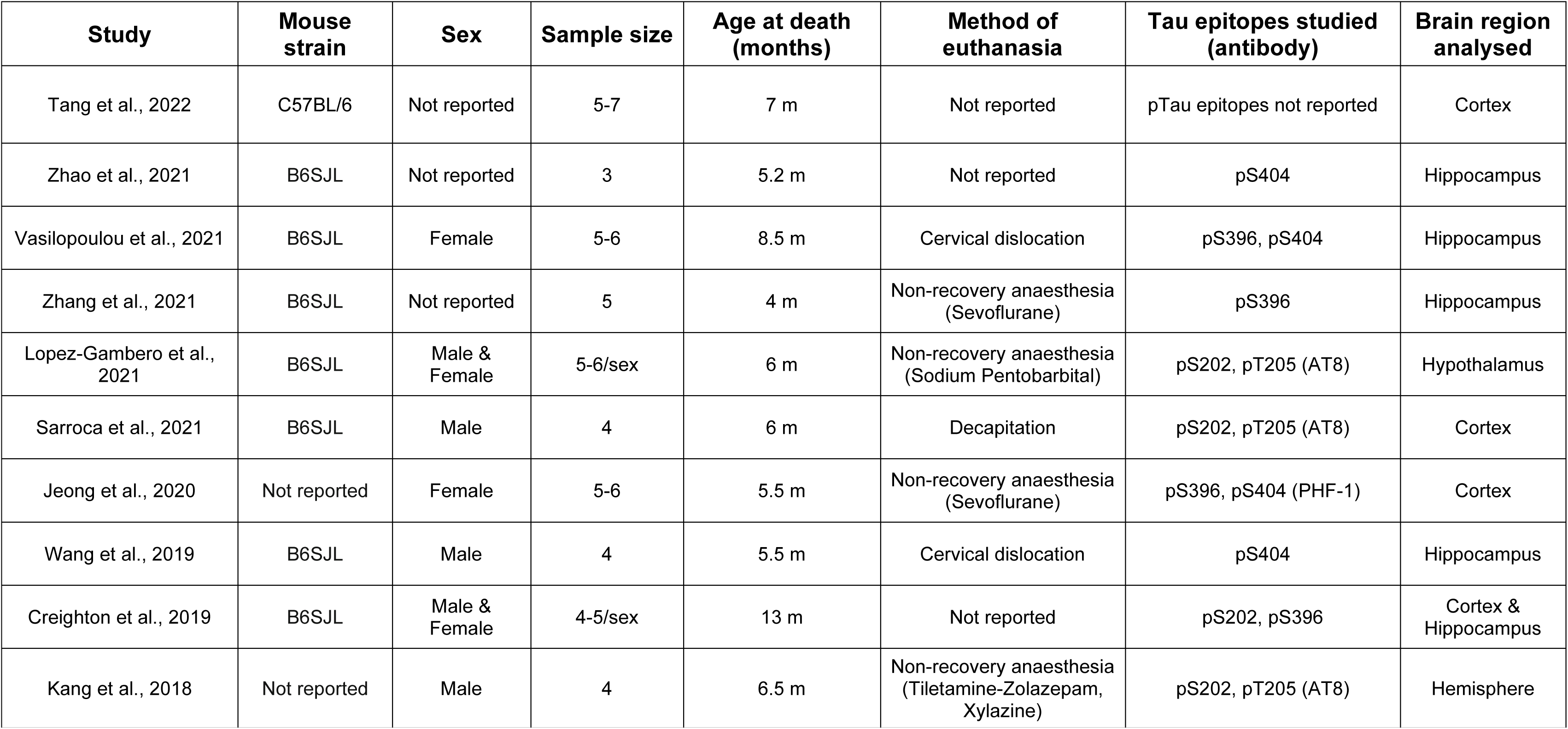
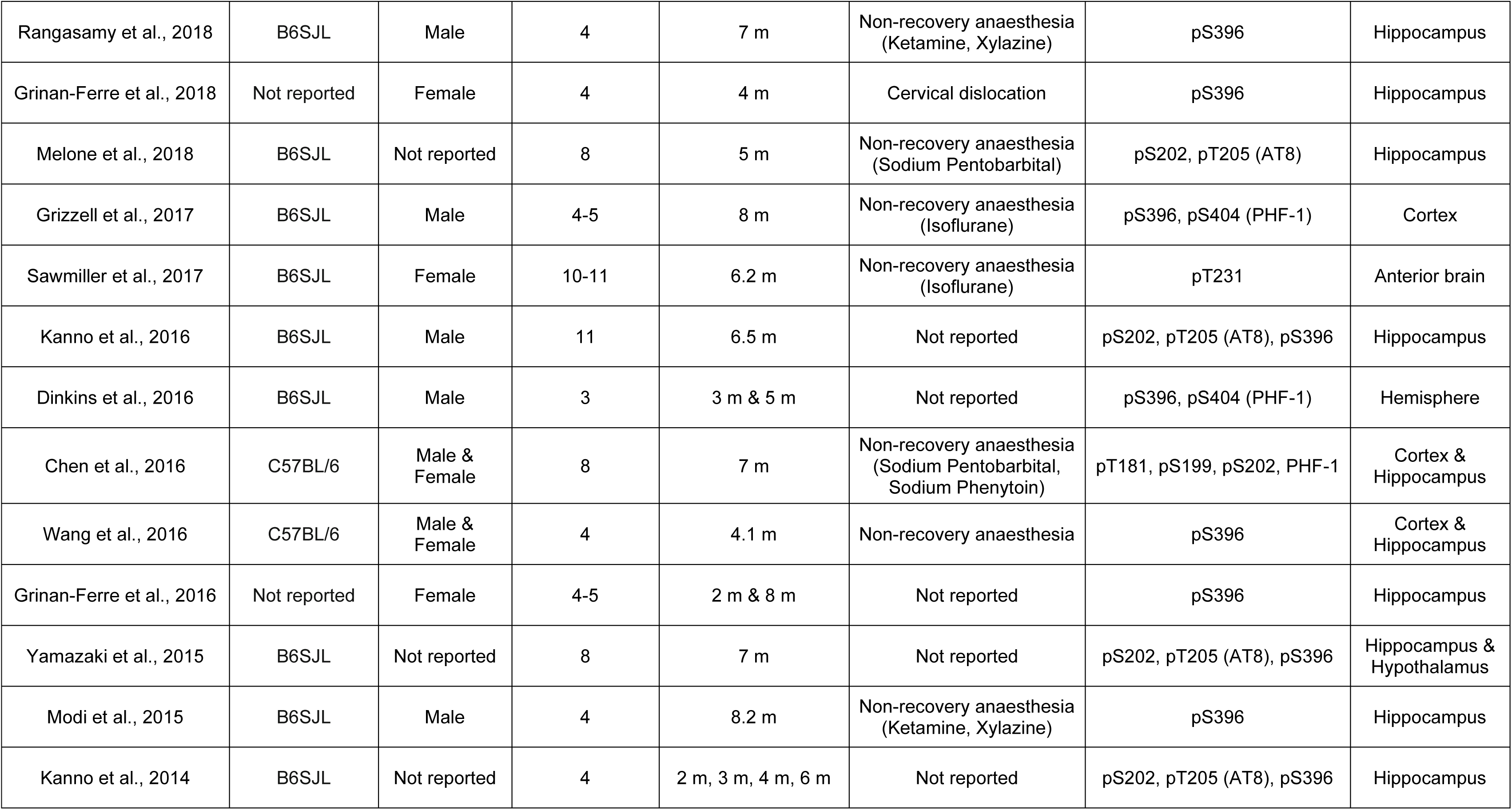
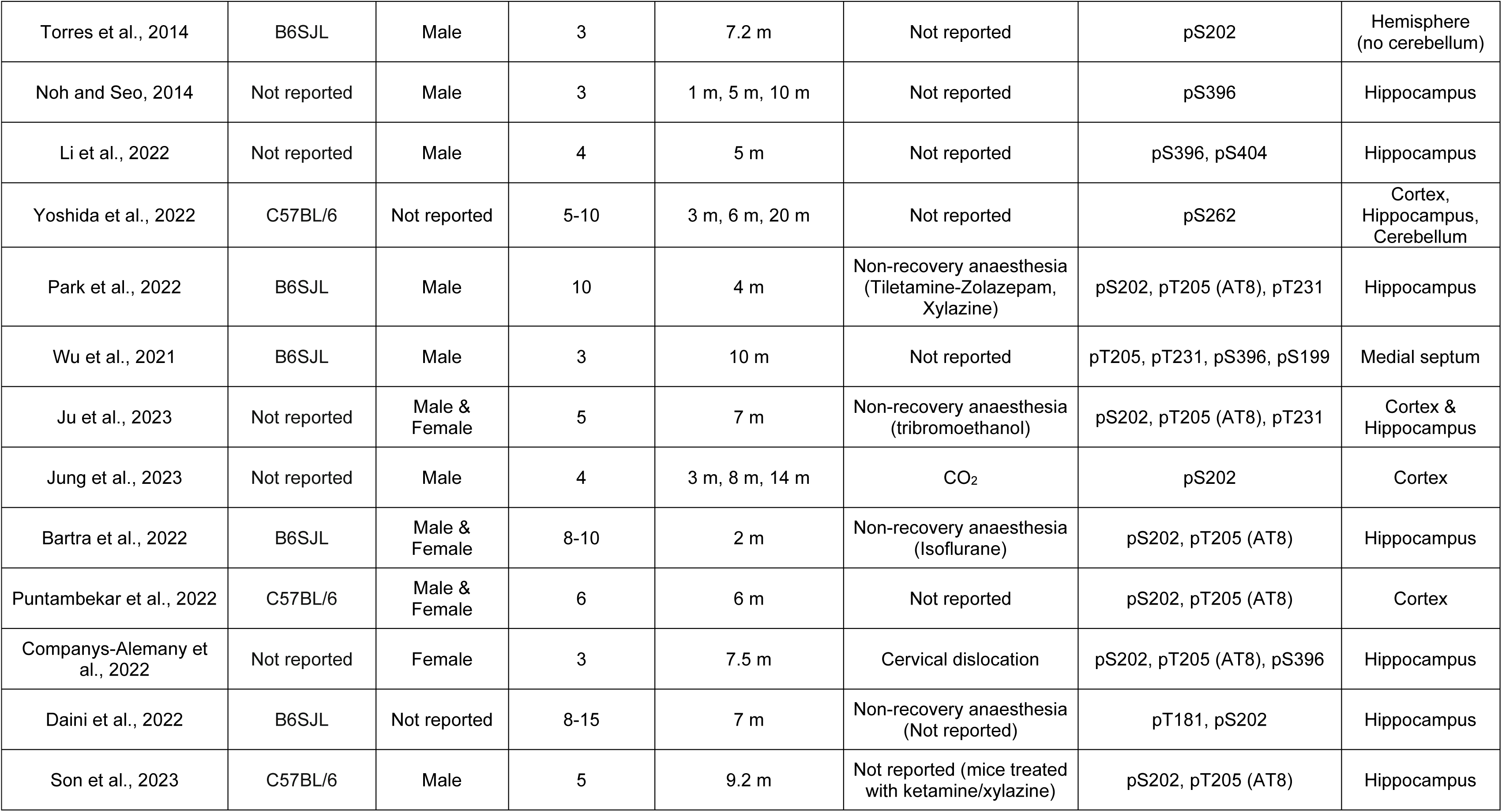
Summary of studies included in the 5xFAD model analysis.

A total of 508 5xFAD and control mice were included in the analysis. The median number of animals per group was 4 (range: 3-11), and the median age was 6 months old (range: 1-20 months). Most studies (87.2%) were conducted using mice younger than 9 months of age. Male and female subjects were used exclusively in 14 (38.9%) and 6 (16.7%) studies, respectively. Both sexes were used in 8 studies (22.2%), while 8 studies did not provide information regarding sex (22.2%). Terminal anaesthesia, using several different agents, was employed as a means of euthanasia in 15 studies (41.7%), cervical dislocation/decapitation in 5 studies (13.9%) and CO_2_ in 1 study (2.8%). The method of euthanasia was not clearly stated in 15 studies (41.7%). Tau phosphorylation was assessed primarily in the cortex and/or the hippocampus (30 studies; 83.3%) or preparations containing these brain regions (4 studies; 11.1%), using antibodies directed to the PR domain (14 studies; 38.9%), the CT region of tau (14 studies; 38.9%), or at both tau regions (7 studies; 19.4%). The exact tau epitopes assessed for phosphorylation were not reported in 1 study (2.8%).

### 3.2 Risk of bias: 5xFAD model

Fig. 2 shows the RoB assessment, which was performed by supplementing the 10 entries included in the SYRCLE tool (Fig. 2A) with 3 questions aiming to assess reporting bias (Fig. 2B). For the majority of items in the SYRCLE tool, most studies received a score of ‘unclear’, as they did not report information that could be used to evaluate selection (entries 1-3), performance (entries 4-5), detection (entries 6-7), attrition (entry 8), reporting (entry 9), and ‘other’ types of bias (entry 10). The lack of information was confirmed by using the additional three questions that were designed to assess the quality of reporting. Most studies did not report any form of randomization, blinding or power calculation in their design.

**Fig. 2.**
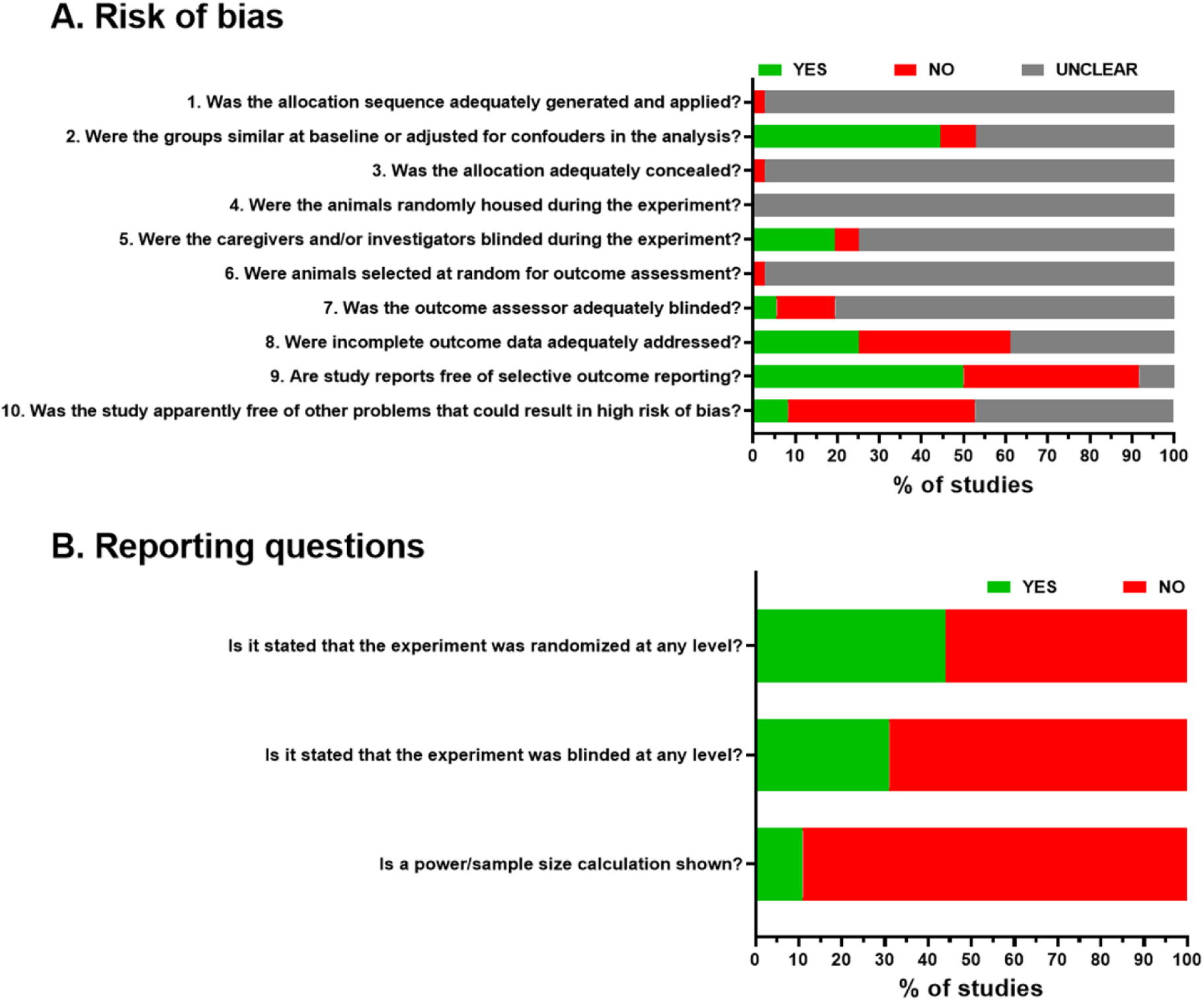
Risk of bias assessment for the studies included in the analysis of the 5xFAD model. Summary bar plots, showing the proportion of studies with a given risk of bias judgement within each domain, obtained by using (A) the SYRCLE tool, and (B) three questions on reporting randomization, blinding and power calculations.

### 3.3 Meta-analysis: 5xFAD model

The 47 comparisons of soluble tau phosphorylation in transgenic vs. control mice (coming out of 36 individual studies) showed a pooled effect size (Hedge’s g ± standard error) of 1.34 ± 0.17 [CI: 0.99-1.68; PI: -0.25-2.93]. The effect direction was positive in 45 out of 47 comparisons, indicating higher levels of tau phosphorylation in the 5xFAD model relative to control mice (Z=7.85, *P*<0.000, two-tailed; Fig. 3). Heterogeneity across studies was substantial (Q=112.75, *P*_Q_<0.000; I^2^=59.20%). Age had no effect as a potential moderator of the relationship between genotype and soluble tau phosphorylation (B=-0.03, β=-0.10, SE=0.04, *P*=0.463; Supplementary Fig. S1).

**Fig. 3.**
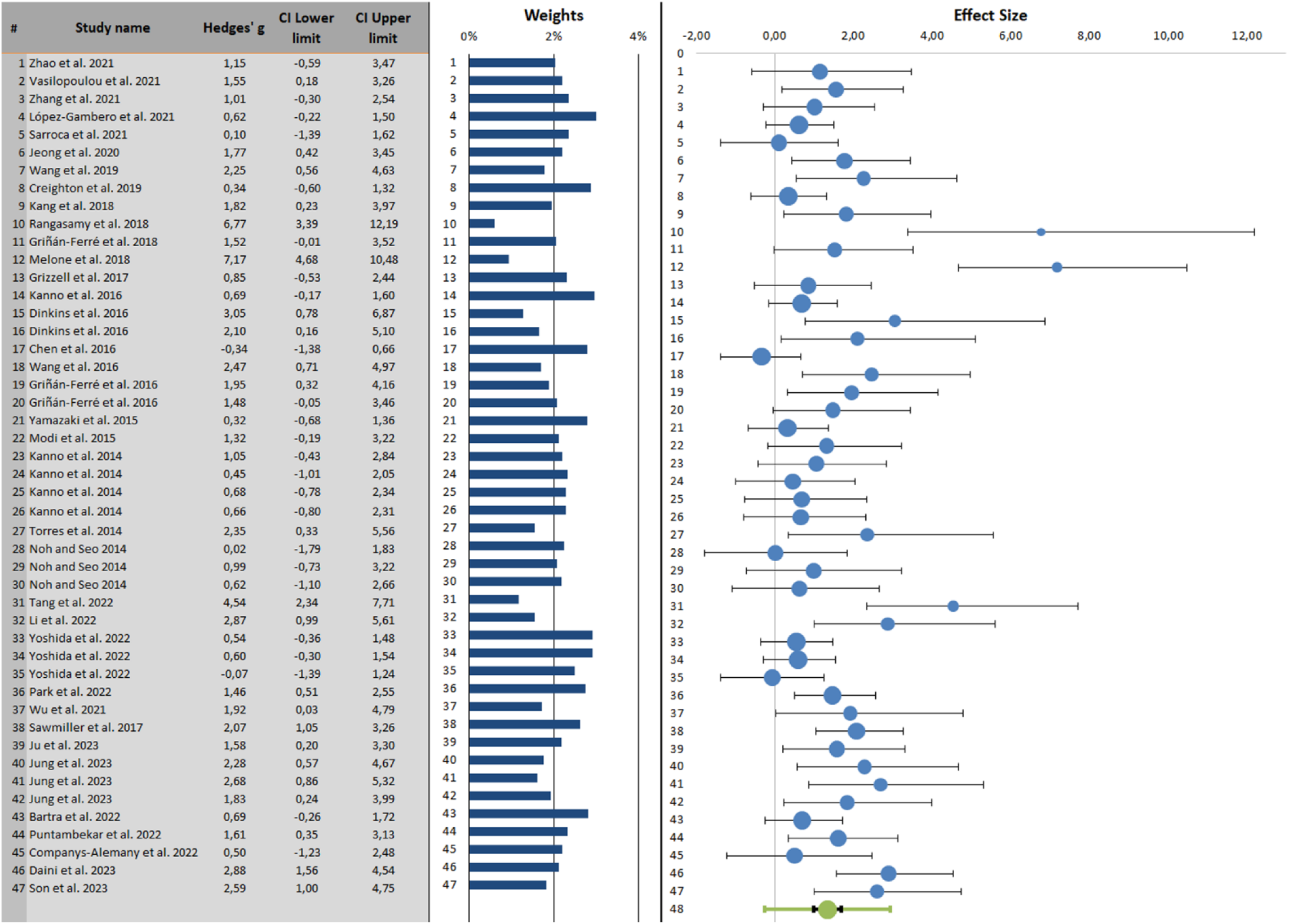
Forest plot of changes in soluble tau phosphorylation in 5xFAD vs. control mice. Point estimates of the effect size ± 95% confidence intervals are shown for each study in blue bullets and horizontal lines, respectively. The relative size of each bullet represents a study’s weight in the generation of the meta-analytic result. The green bullet represents the combined effect size (weighted average effect). The smaller, black interval on each side of the combined effect size is a confidence interval, while the larger, green interval is the prediction interval. The direction of effect size was positive in 45 out of 47 comparisons, indicating significant effects of genotype on the phosphorylation of tau.

Separate, pre-specified meta-analyses were conducted to study the effects of genotype on tau phosphorylation within both the PR and the CT domains of tau. Subgroup analysis examined the effects of brain area (pre-specified subgroup), sex and mouse genetic background (*post-hoc* defined subgroups) on the phosphorylation of the CT and PR domains of soluble tau.

For the CT domain (Fig. 4), there was a pooled effect size of 1.48 ± 0.22 [CI: 1.03-1.94; PI: - 0.19-3.16] and substantial heterogeneity (Q=61.04, *P*_Q_<0.000; I^2^=55.77%). The effect direction was positive in 27 out of 28 comparisons [Z=6.69, *P*<0.000, two-tailed value]. Subgroup analysis showed no differences in CT phosphorylated tau between brain regions (cortex: n=5, g=0.75±0.40; CI: -0.06-1.56; PI: -1.49-2.99; I^2^=59.73%; hippocampus: n=23, g=1.45±0.28; CI: 0.90-2.01; PI: -0.58-3.49; I^2^=64.85%; between group difference: *P*=0.141), and between male and female mice (male: n=13, g=1.61±0.36; CI: 0.90-2.32; PI: -0.33-3.56; I^2^=54.11%; female: n=7, g=1.19±0.30; CI: 0.58-1.80; PI:-0.18-2.56; I^2^=32.38%; between group difference: *P*=0.345). The limited number of studies in congenic mice (n=2) prevented us from comparing CT phosphorylated tau between congenic and hybrid animals (hybrid: n=17; g=1.73±0.32; CI: 1.09-2.36; PI: -0.26-3.71; I^2^=60.40%). Moderator analysis showed that age tended to negatively predict the effect of genotype on the CT phosphorylation of tau (B=-0.12, β=-0.30, SE=0.07, *P*=0.08). With ageing, the differences in tau phosphorylation between transgenic and wild-type animals tended to become less pronounced (Fig. 5A). Age negatively moderated the relationship between genotype and CT tau phosphorylation in hybrid mice (B=-0.20, β=-0.45, SE=0.09, *P*=0.027; Fig. 5B), female mice (B=-0.20, β=-0.87, SE=0.08, *P*=0.009; Fig. 5C) and the cortex (B=-0.22, β=-0.88, SE=0.08, *P*=0.005; Fig. 5D).

**Fig. 4.**
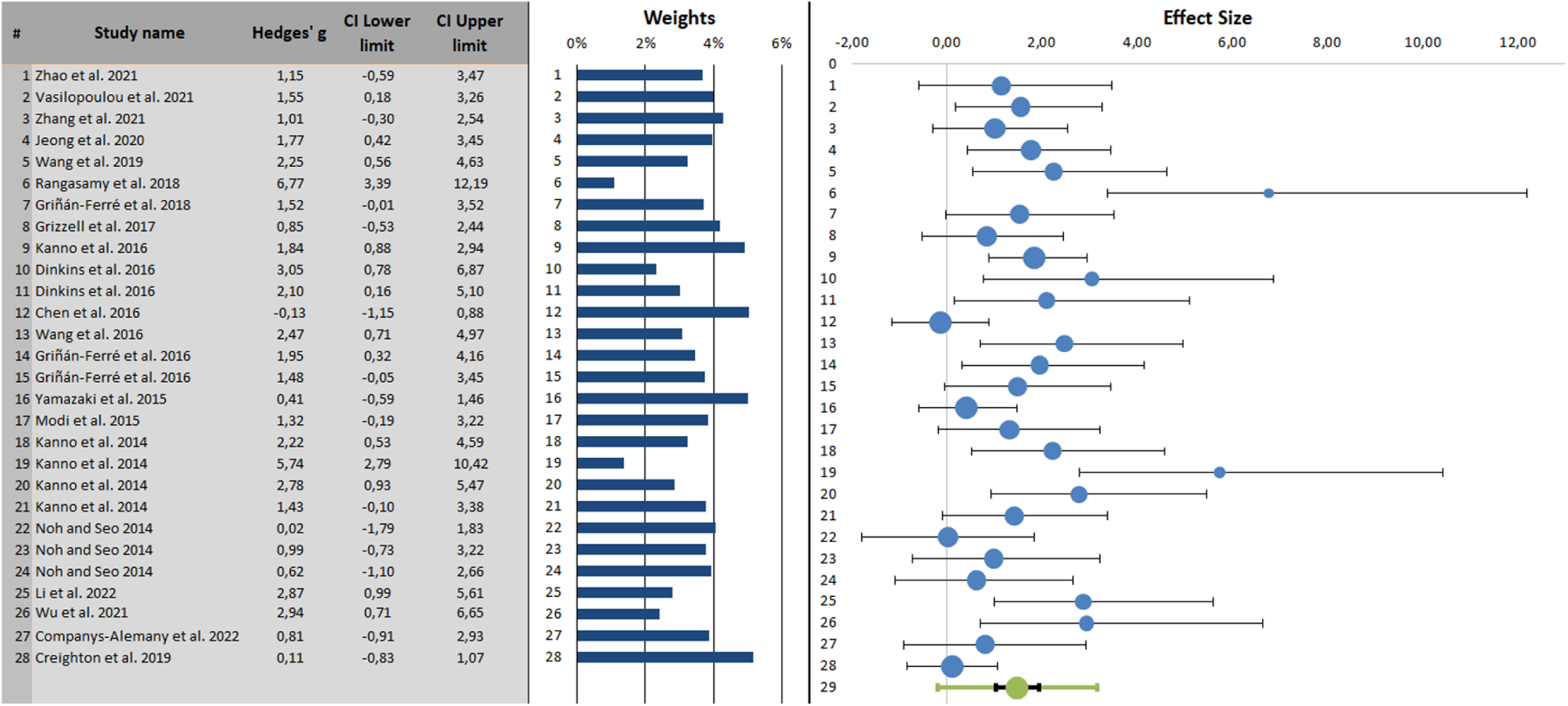
Forest plot of changes in C-terminal domain phosphorylated tau in 5xFAD vs. control mice. Point estimates of the effect size ± 95% confidence intervals are shown for each study in blue bullets and horizontal lines, respectively. The relative size of each bullet represents a study’s weight in the generation of the meta-analytic result. The green bullet represents the combined effect size (weighted average effect), along with the confidence interval (black horizontal line) and the prediction interval (green horizontal line). The direction of effect size was positive in 27 out of 28 comparisons, indicating significant effects of genotype on the phosphorylation of tau at the C-terminal domain.

**Fig. 5.**
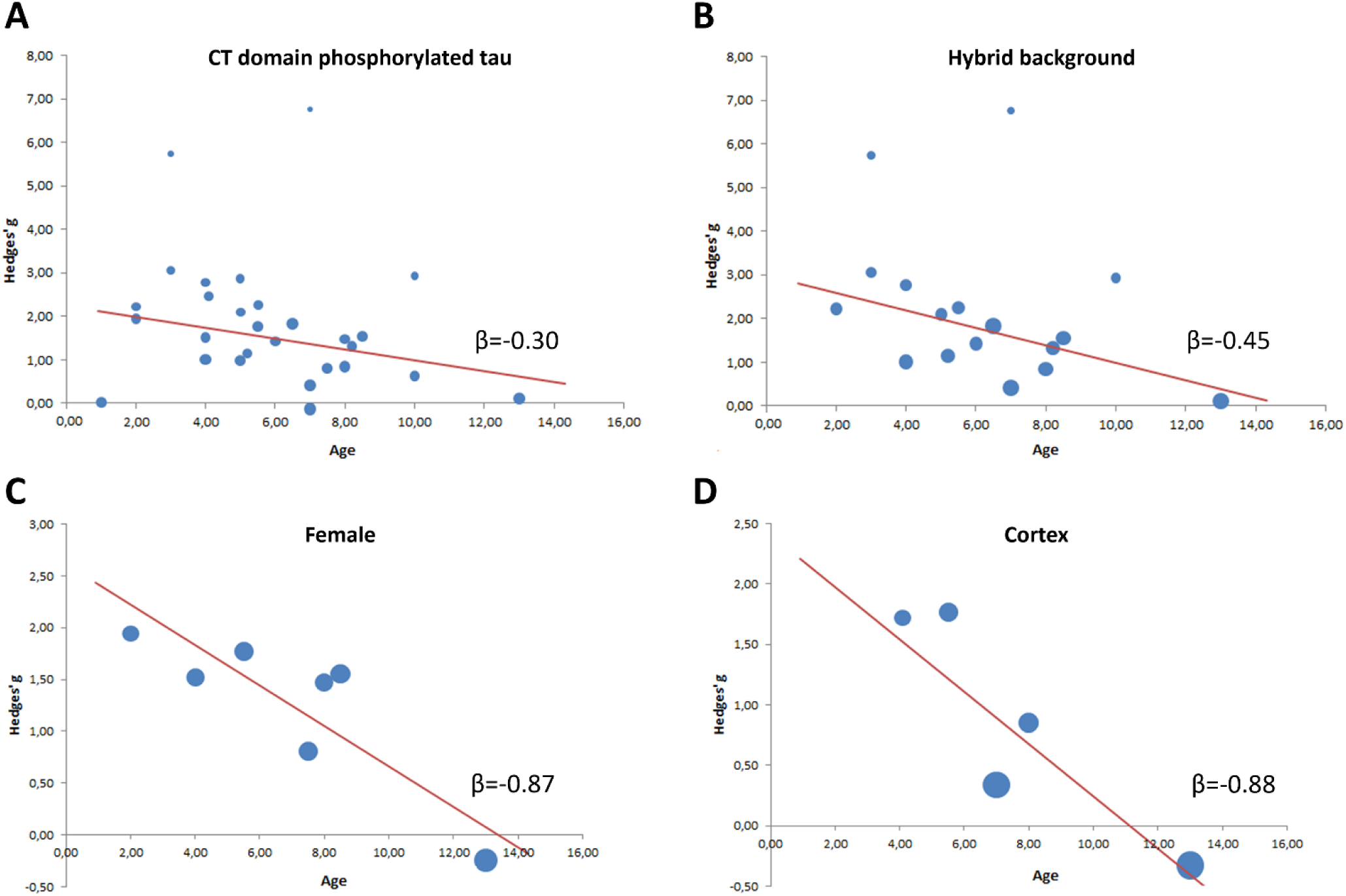
Meta-regression performed with age as moderator in 5xFAD vs. control mice. Scatter plots and regression lines for the moderating effects of age on the phosphorylation of tau at the CT domain, for (A) all retrieved studies, (B) studies using hybrid mice, (C) female mice, and (D) studies using cortical preparations. Each data point represents the effect size observed at a specific time-point. In all cases, there was a progressive decrease in the magnitude of hyperphosphorylation in transgenic mice with ageing.

For the PR domain of tau (Fig. 6), there was a pooled effect size of 1.02 ± 0.24 [CI: 0.53-1.52; PI: -0.75-2.80] and substantial heterogeneity (Q=80.49, *P*_Q_<0.000; I^2^=67.70%). The effect direction was positive in 23 out of 27 comparisons [Z=4.27, *P*<0.000, two-tailed value]. Subgroup analysis showed no differences in phosphorylation at the PR domain of tau between brain regions (cortex: n=11, g=0.89±0.34; CI: 0.22-1.56; PI: -1.19-2.97; I^2^=69.10%; hippocampus: n=18, g=0.76±0.33; CI: 0.10-1.41; PI: -1.26-2.77; I^2^=72.69%; between group difference: *P*=0.750), and between mice of different genetic background (hybrid: n=15; g=0.98±0.37; CI: 0.25-1.71; PI: -1.12-3.08; I^2^=73.00%; congenic: n=5; g=0.54±0.42; CI: -0.31-1.39; PI: -1.54-2.62; I^2^=58.76%; between group difference: *P*=0.338). Subgroup comparisons between male (n=11, g=1.33±0.29; CI: 0.76-1.91; PI: -0.39-3.06; I^2^=53.32) and female mice were not performed, due to the small number of studies using female animals (n=4). Age did not moderate the relationship between PR phosphorylated tau and genotype (B=-0.01, β=-0.03, SE=0.05, *P*=0.848; Supplementary Fig. S2.A), in any of the subgroups examined (Supplementary Fig. S2.B-G).

**Fig. 6.**
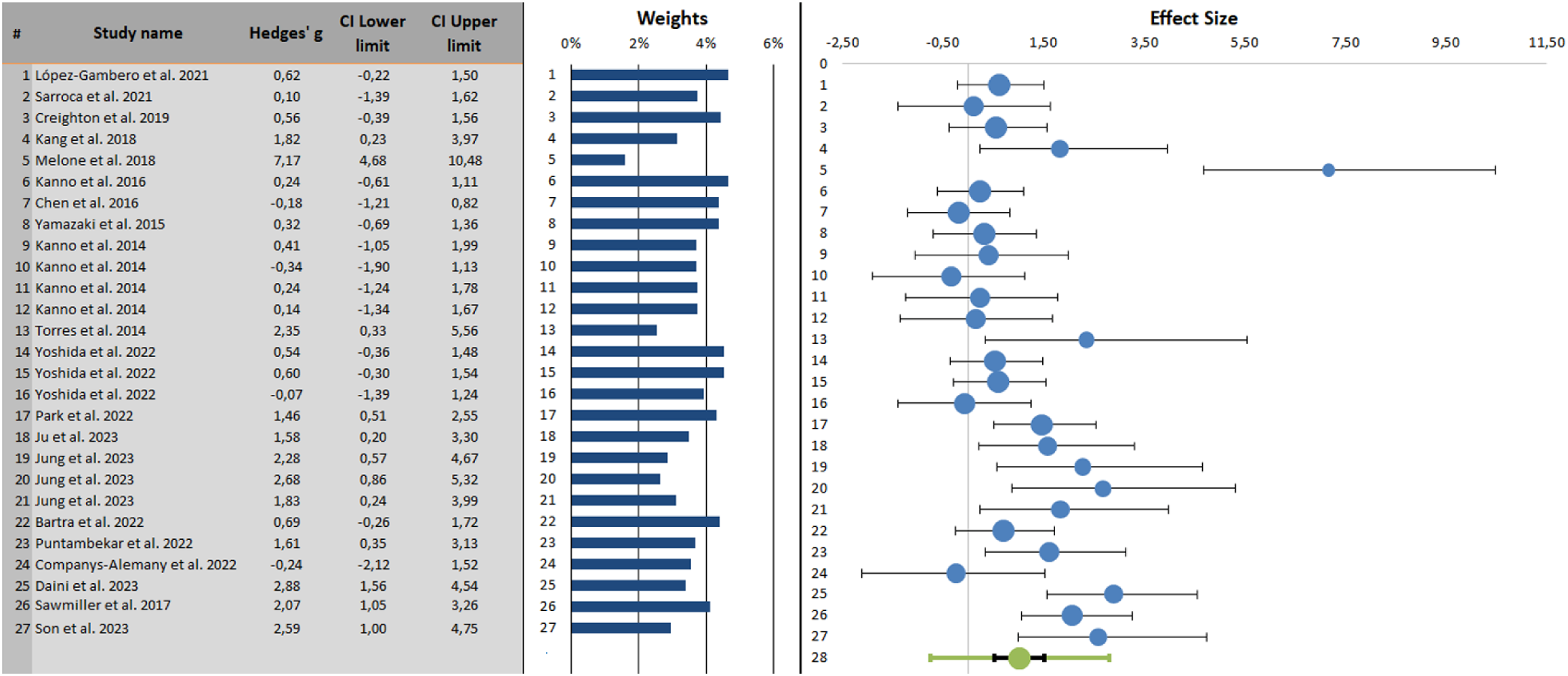
Forest plot of changes in proline-rich phosphorylated tau in 5xFAD vs. control mice. Point estimates of the effect size ± 95% confidence intervals are shown for each study in blue bullets and horizontal lines, respectively. The relative size of each bullet represents a study’s weight in the generation of the meta-analytic result. The green bullet represents the combined effect size (weighted average effect), along with the confidence interval (black horizontal line) and the prediction interval (green horizontal line). The direction of effect size was positive in 23 out of 27 comparisons, indicating significant effects of genotype on the phosphorylation of tau at the proline-rich domain.

### 3.4 Publication bias: 5xFAD model

Egger’s regression analysis revealed funnel plot asymmetry for the studies measuring total soluble tau (intercept: 7.08±0.65, t=10.86, *P*<0.000), CT-phosphorylated tau (intercept: 6.44±0.58, t=11.02, *P*<0.000) and PR-phosphorylated tau (intercept: 8.57±1.39, t=6.19, *P*<0.000). Trim & fill analysis using leftmost/rightmost estimators imputed 8, 5 and 10 missing studies for total tau, CT-phosphorylated, and PR-phosphorylated tau, respectively. Funnel plots and estimates of adjusted combined effect sizes are shown in Fig. 7A-C.

**Fig. 7.**
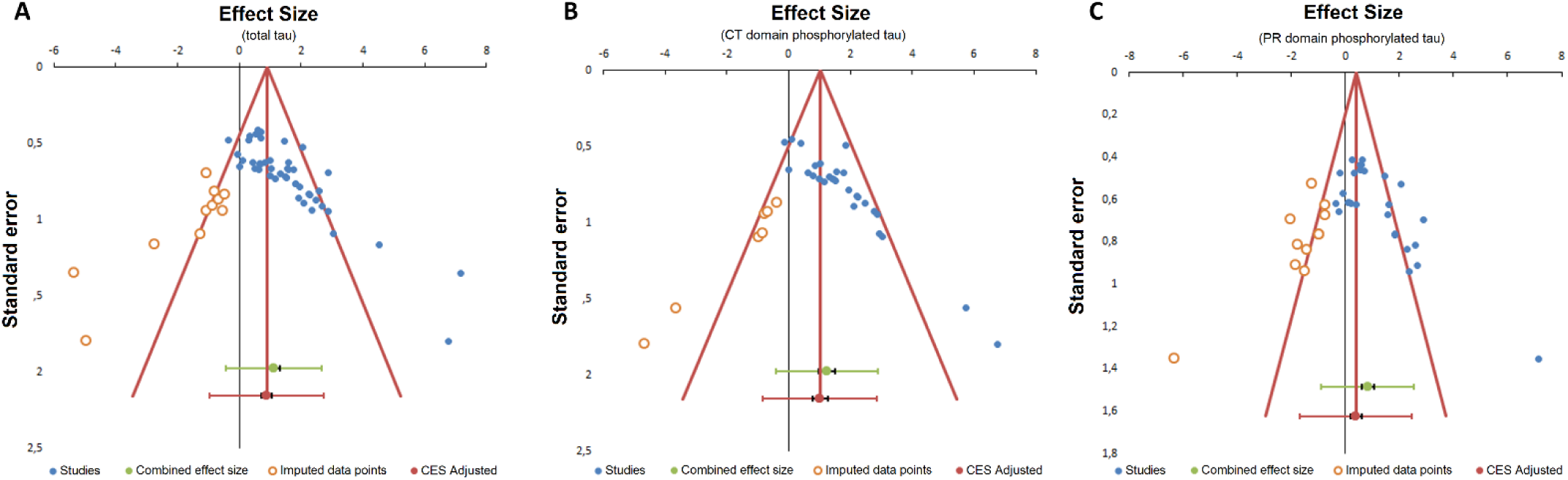
Funnel plot of meta-analysis examining soluble tau phosphorylation in 5xFAD vs. control mice. (A) total soluble tau phosphorylation (B) tau phosphorylated at the C-terminal domain (C) tau phosphorylated at the proline-rich domain. Each symbol represents an independent comparison. Adjusted combined effect sizes (CES) were: (A) 0.92±0.21 (CI: 0.50-1.34; PI: -0.92-2.76); (B): 1.17±0.26 (CI: 0.65-1.69; PI: -0.68-3.02); (C): 0.06±0.23 (CI: -0.41-0.52; PI: -1.94-2.05).

### 3.5 Study selection and characteristics: *APP*_swe_/*PSEN1*_dE9_ model

The flowchart of study selection for the *APP*_swe_/*PSEN1*_dE9_ model is presented in Fig. 8. Searching in PubMed and Scopus retrieved 567 articles, of which 91 were duplicates between and within databases. After screening the remaining 476 records, a total of 66 primary research articles were assessed for eligibility. Of them, 19 articles did not measure tau phosphorylation compared to control mice, 10 articles reported tau phosphorylation exclusively in a qualitative manner (no quantification), 6 articles quantified tau using methods other than immunoblotting or ELISA, and 4 articles lacked vital study information. The characteristics of the 27 articles that were included in the study are shown in Table 2.

**Fig. 8.**
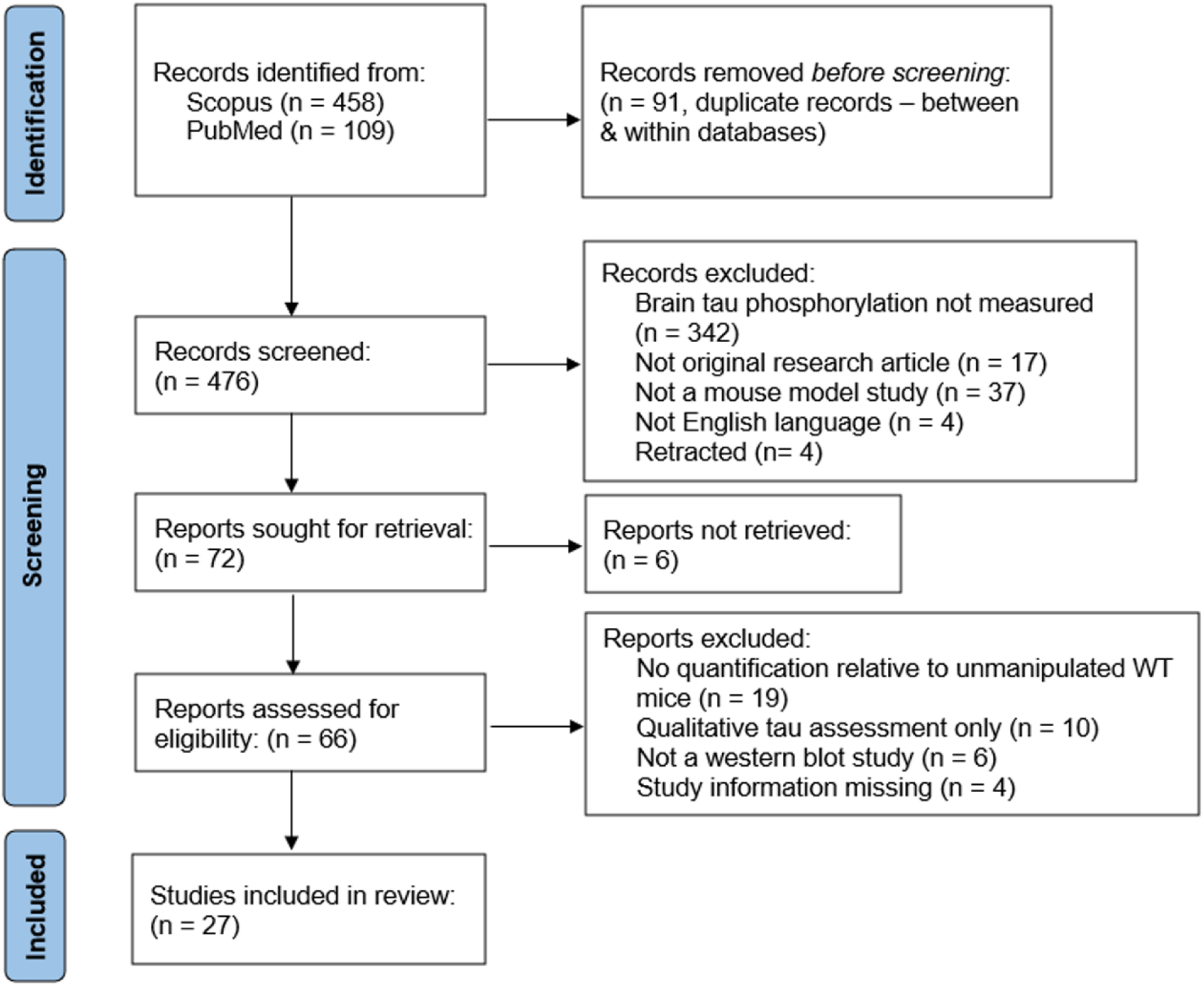
Flowchart of study search and selection for the *APP*_swe_/*PSEN1*_dE9_ model.

**Table 2.**
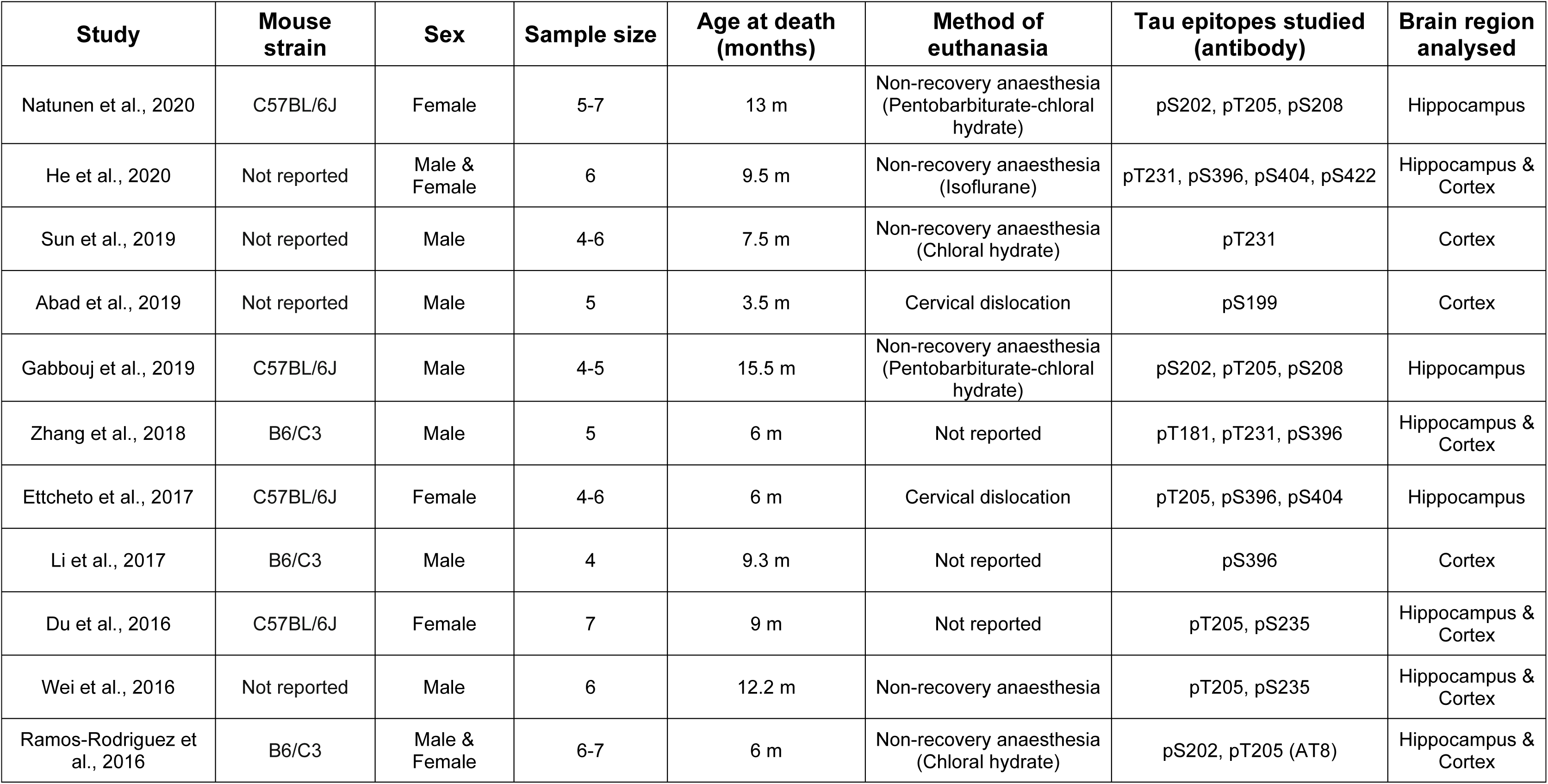
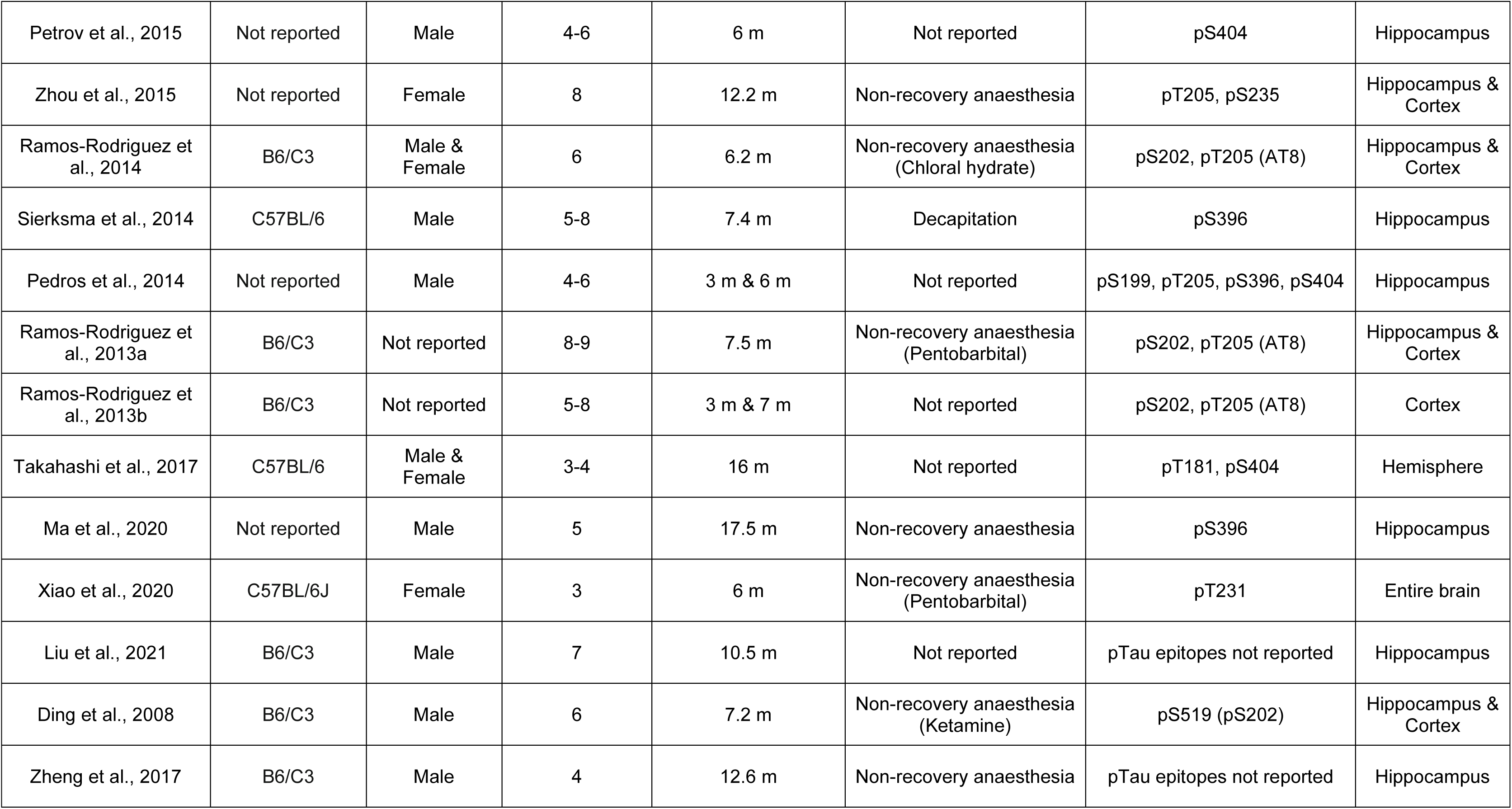
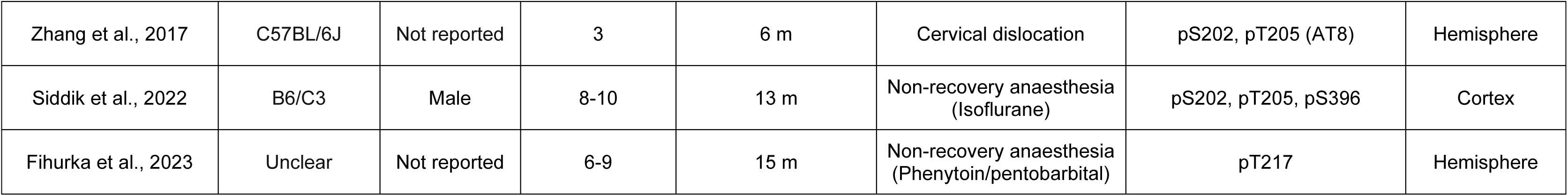
Summary of studies included in the analysis of the APPswe/PSEN1dE9 model.

| **Study** | **Mouse strain** | **Sex** | **Sample size** | **Age at death (months)** | **Method of euthanasia** | **Tau epitopes studied (antibody)** | **Brain region analysed** |
| --- | --- | --- | --- | --- | --- | --- | --- |
| Natunen et al., 2020 | C57BL/6J | Female | 5-7 | 13 m | Non-recovery anaesthesia (Pentobarbiturate-chloral hydrate) | pS202, pT205, pS208 | Hippocampus |
| He et al., 2020 | Not reported | Male & Female | 6 | 9.5 m | Non-recovery anaesthesia (Isoflurane) | pT231, pS396, pS404, pS422 | Hippocampus & Cortex |
| Sun et al., 2019 | Not reported | Male | 4-6 | 7.5 m | Non-recovery anaesthesia (Chloral hydrate) | pT231 | Cortex |
| Abad et al., 2019 | Not reported | Male | 5 | 3.5 m | Cervical dislocation | pS199 | Cortex |
| Gabbouj et al., 2019 | C57BL/6J | Male | 4-5 | 15.5 m | Non-recovery anaesthesia (Pentobarbiturate-chloral hydrate) | pS202, pT205, pS208 | Hippocampus |
| Zhang et al., 2018 | B6/C3 | Male | 5 | 6 m | Not reported | pT181, pT231, pS396 | Hippocampus & Cortex |
| Ettcheto et al., 2017 | C57BL/6J | Female | 4-6 | 6 m | Cervical dislocation | pT205, pS396, pS404 | Hippocampus |
| Li et al., 2017 | B6/C3 | Male | 4 | 9.3 m | Not reported | pS396 | Cortex |
| Du et al., 2016 | C57BL/6J | Female | 7 | 9 m | Not reported | pT205, pS235 | Hippocampus & Cortex |
| Wei et al., 2016 | Not reported | Male | 6 | 12.2 m | Non-recovery anaesthesia | pT205, pS235 | Hippocampus & Cortex |
| Ramos-Rodriguez et al., 2016 | B6/C3 | Male & Female | 6-7 | 6 m | Non-recovery anaesthesia (Chloral hydrate) | pS202, pT205 (AT8) | Hippocampus & Cortex |
| Petrov et al., 2015 | Not reported | Male | 4-6 | 6 m | Not reported | pS404 | Hippocampus |
| Zhou et al., 2015 | Not reported | Female | 8 | 12.2 m | Non-recovery anaesthesia | pT205, pS235 | Hippocampus & Cortex |
| Ramos-Rodriguez et al., 2014 | B6/C3 | Male & Female | 6 | 6.2 m | Non-recovery anaesthesia (Chloral hydrate) | pS202, pT205 (AT8) | Hippocampus & Cortex |
| Sierksma et al., 2014 | C57BL/6 | Male | 5-8 | 7.4 m | Decapitation | pS396 | Hippocampus |
| Pedros et al., 2014 | Not reported | Male | 4-6 | 3 m & 6 m | Not reported | pS199, pT205, pS396, pS404 | Hippocampus |
| Ramos-Rodriguez et al., 2013a | B6/C3 | Not reported | 8-9 | 7.5 m | Non-recovery anaesthesia (Pentobarbital) | pS202, pT205 (AT8) | Hippocampus & Cortex |
| Ramos-Rodriguez et al., 2013b | B6/C3 | Not reported | 5-8 | 3 m & 7 m | Not reported | pS202, pT205 (AT8) | Cortex |
| Takahashi et al., 2017 | C57BL/6 | Male & Female | 3-4 | 16 m | Not reported | pT181, pS404 | Hemisphere |
| Ma et al., 2020 | Not reported | Male | 5 | 17.5 m | Non-recovery anaesthesia | pS396 | Hippocampus |
| Xiao et al., 2020 | C57BL/6J | Female | 3 | 6 m | Non-recovery anaesthesia (Pentobarbital) | pT231 | Entire brain |
| Liu et al., 2021 | B6/C3 | Male | 7 | 10.5 m | Not reported | pTau epitopes not reported | Hippocampus |
| Ding et al., 2008 | B6/C3 | Male | 6 | 7.2 m | Non-recovery anaesthesia (Ketamine) | pS519 (pS202) | Hippocampus & Cortex |
| Zheng et al., 2017 | B6/C3 | Male | 4 | 12.6 m | Non-recovery anaesthesia | pTau epitopes not reported | Hippocampus |
| Zhang et al., 2017 | C57BL/6J | Not reported | 3 | 6 m | Cervical dislocation | pS202, pT205 (AT8) | Hemisphere |
| Siddik et al., 2022 | B6/C3 | Male | 8-10 | 13 m | Non-recovery anaesthesia (Isoflurane) | pS202, pT205, pS396 | Cortex |
| Fihurka et al., 2023 | Unclear | Not reported | 6-9 | 15 m | Non-recovery anaesthesia (Phenytoin/pentobarbital) | pT217 | Hemisphere |

A total of 316 transgenic and wild-type mice were included in the analysis. The median number of animals per genotype was 5 (range: 3-11), and the median age 7.5 months old (range: 3-17.5 months). Male and female subjects were used exclusively in 14 (51.9%) and 5 (18.5%) studies, respectively. Both sexes were used in 4 studies (14.8%), while 4 studies did not provide information regarding sex (14.8%). Terminal anaesthesia was employed as a means of euthanasia in 15 studies (55.6%), cervical dislocation/decapitation in 4 studies (14.8%), while the method of euthanasia was not reported in 8 studies (29.6%). In all studies, tau phosphorylation was assessed in preparations containing the cortex and/or the hippocampus, using antibodies directed to the PR (15 studies, 55.6%) and the CT domain of tau (4 studies, 14.8%), or at both tau regions (6 studies; 22.2%). The exact tau epitopes assessed for phosphorylation were not reported in 2 studies (7.4%).

### 3.6 Risk of bias: *APP*_swe_/*PSEN1*_dE9_ model

The results from the RoB assessment, obtained using the SYRCLE tool along with three additional questions assessing reporting bias, are shown in Fig. 9A & Fig. 9B, respectively. As with the 5xFAD mice, most studies were evaluated to have high risk of bias.

**Fig. 9.**
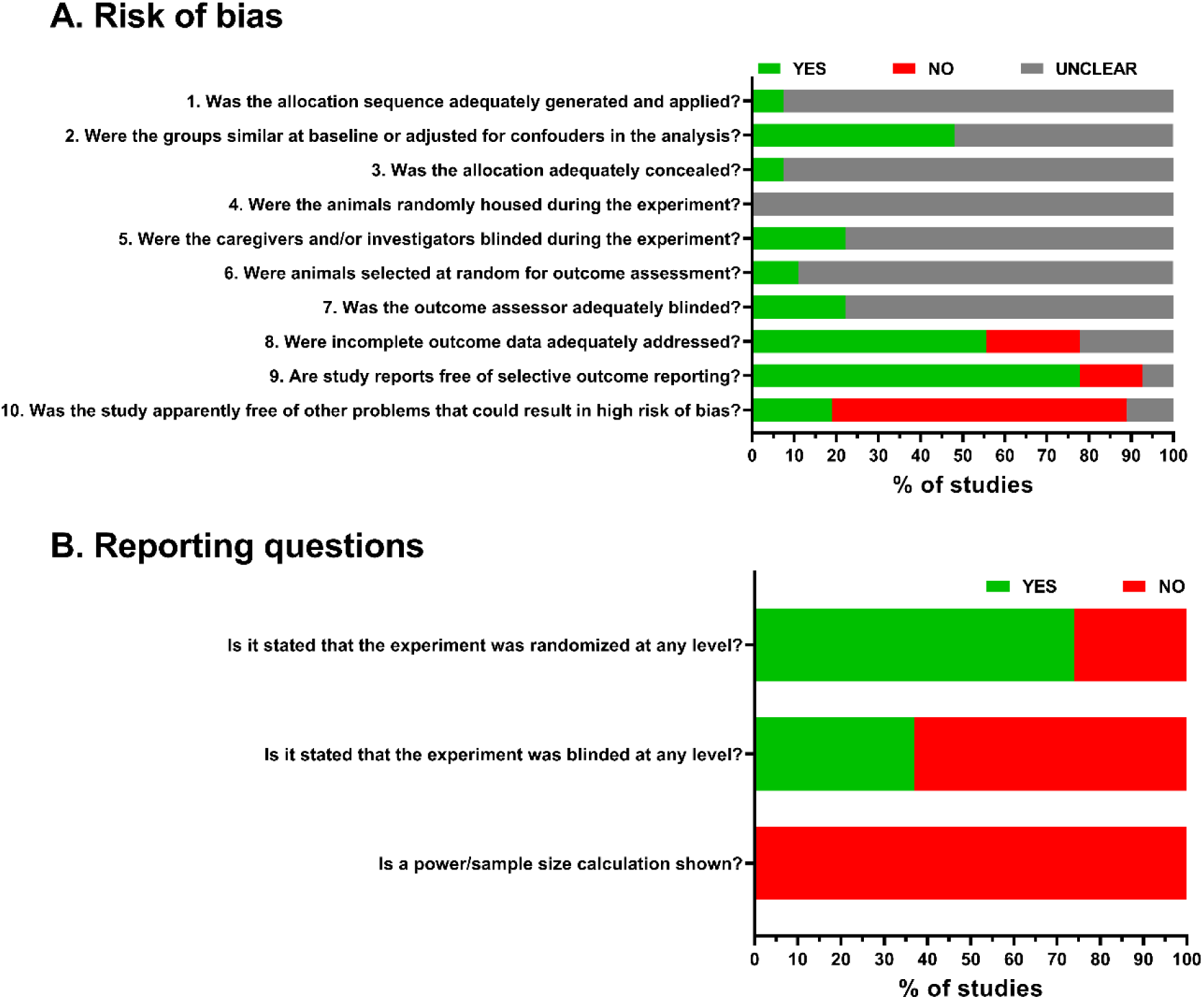
Risk of bias assessment for the studies included in the analysis of the *APP*_swe_/*PSEN1*_dE9_ model. Summary bar plots, showing the proportion of studies with a given risk of bias judgement within each domain, obtained by using (A) the SYRCLE tool, and (B) three questions on reporting randomization, blinding and power calculations.

### 3.7 Meta-analysis: *APP*_swe_/*PSEN1*_dE9_ model

The 29 comparisons of soluble tau phosphorylation in transgenic vs. control mice (coming out of 27 individual studies) showed a pooled effect size of 1.44 ± 0.34 [CI: 0.73-2.15; PI: -1.23- 4.11]. The effect direction was positive in 23 out of 29 comparisons, indicating higher levels of tau phosphorylation in *APP*_swe_/*PSEN1*_dE9_, relative to control mice (Z=4.18, P<0.000, two-tailed; Fig. 10). Heterogeneity was substantial (Q=126.57, *P*_Q_<0.000; I^2^=77.88%). Age had no effect as a potential moderator of the relationship between genotype and soluble tau phosphorylation (B=0.02, β=0.04, SE=0.07, *P*=0.784; Supplementary Fig. S3).

**Fig. 10.**
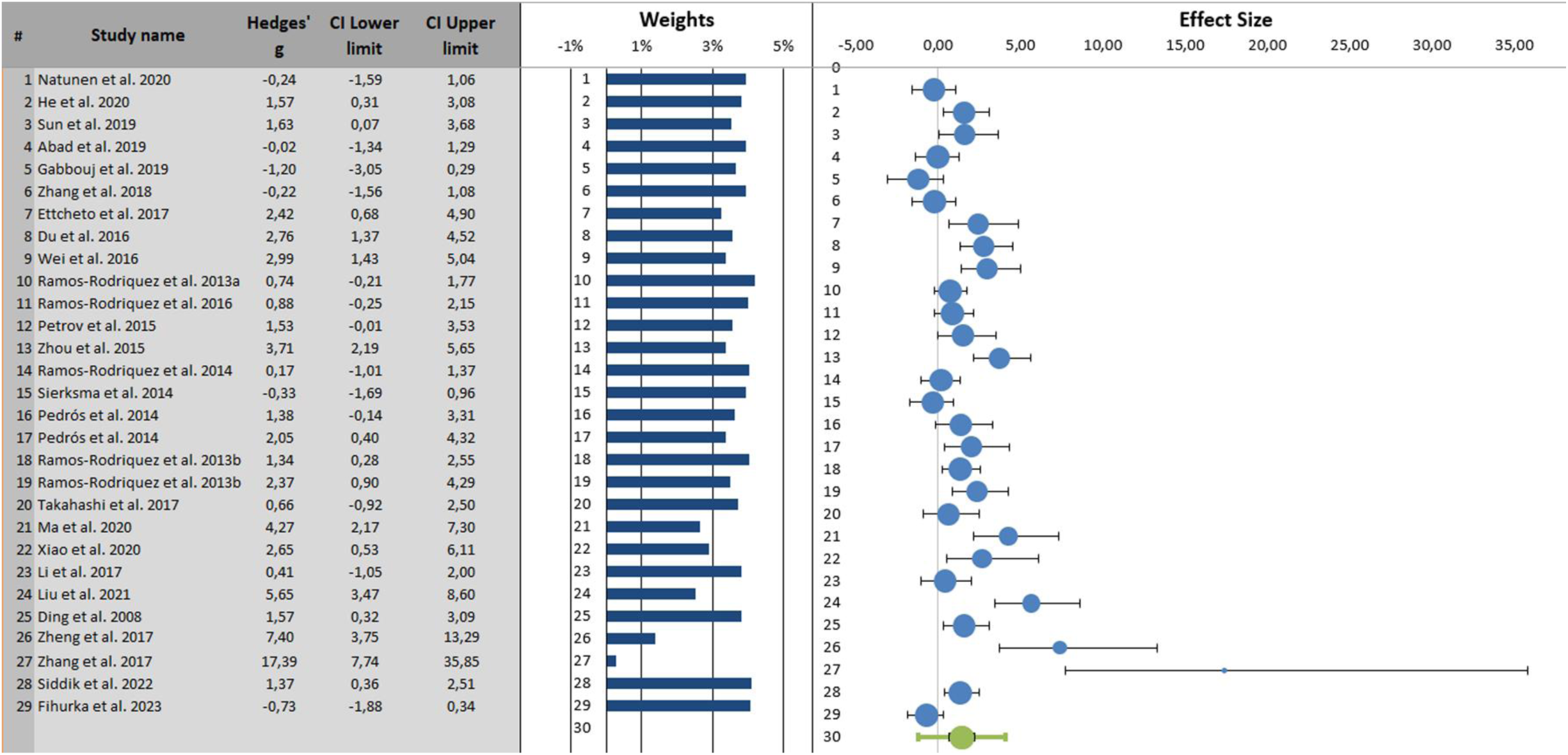
Forest plot of changes in soluble tau phosphorylation in *APP*_swe_/*PSEN1*_dE9_ vs. control mice. Point estimates of the effect size ± 95% confidence intervals are shown for each study in blue bullets and horizontal lines, respectively. The relative size of each bullet represents a study’s weight in the generation of the meta-analytic result. The green bullet represents the combined effect size (weighted average effect). The smaller, black interval on each side of the combined effect size is a confidence interval, while the larger, green interval is the prediction interval. The direction of effect size was positive in 23 out of 29 comparisons, indicating significant effects of genotype on the phosphorylation of soluble tau.

For the CT domain of tau (Fig. 11), there was a pooled effect size of 1.18 ± 0.37 [CI: 0.38-1.99; PI: -1.06-3.43] and substantial heterogeneity across studies (Q=33.30, *P*_Q_<0.000; I^2^=66.96%). The effect direction was positive in 10 out of 12 comparisons [Z=3.24, *P*=0.001, two-tailed value]. Age had no influence on the relationship between genotype and the CT phosphorylation of tau (B=0.05, β=0.18, SE=0.09, *P*=0.541; Supplementary Fig. S4.A). The number of studies was too small to conduct subgroup comparisons across brain regions (cortex n=4; hippocampus n=8), sexes (male n=11; female n=2) and backgrounds (congenic n=3; hybrid n=5). In male mice (g=0.97±0.41; CI: 0.04-1.89; PI: -1.52-3.45; I^2^=71.07%), age did not significantly moderate the relationship between genotype and CT tau phosphorylation (B=0.16, β=0.43, SE=0.11, *P*=0.143; Supplementary Fig. S4.B). Similar results were obtained in the hippocampus (g=1.45±0.49; CI: 0.30-2.60; PI: -1.35-4.26; I^2^=69.93%; age: B=0.20, β=0.58, SE=0.12, *P*=0.09; Supplementary Fig. S4.C). For the PR domain of tau (Fig. 12), there was a pooled effect size of 1.21 ± 0.33 [CI: 0.54-1.89; PI: -1.22-3.65] and substantial heterogeneity (Q=80.49, *P*_Q_<0.000; I^2^=75.31%). The effect direction was positive in 18 out of 23 comparisons [Z=3.73, *P*<0.000, two-tailed value]. Age positively moderated the relationship between genotype and PR tau phosphorylation in the cortex (B=0.29, β=0.70, SE=0.09, *P*=0.001; Fig. 13), but not in any other groups analyzed (Supplementary Fig. S5.A-F). Subgroup analysis showed no differences in phosphorylation at the PR domain of tau between brain regions (cortex: n=14, g=1.65±0.35; CI: 0.95-2.35; PI: - 0.70-4.00; I^2^=72.61%; hippocampus: n=14, g=1.12±0.47; CI: 0.20-2.04; PI: -2.02-4.26; I^2^=81.89%; between group difference: *P*=0.311), and between mice of different sex (male: n=13, g=1.19±0.50; CI: 0.19-2.18; PI: -1.71-4.08; I^2^=74.94%; female: n=8, g=1.56±0.57; CI: 0.43-2.69; PI:-2.00-5.11; I^2^=77.99%; between group difference: *P*=0.594) and genetic background (hybrid: n=8; g=1.07±0.32; CI: 0.43-1.71; PI: -0.70-2.84; I^2^=58.92%; congenic: n=7; g=1.10±1.07; CI: -1.03-3.23; PI: -3.61-5.81; I^2^=81.93%; between group difference: *P*=0.971).

**Fig. 11.**
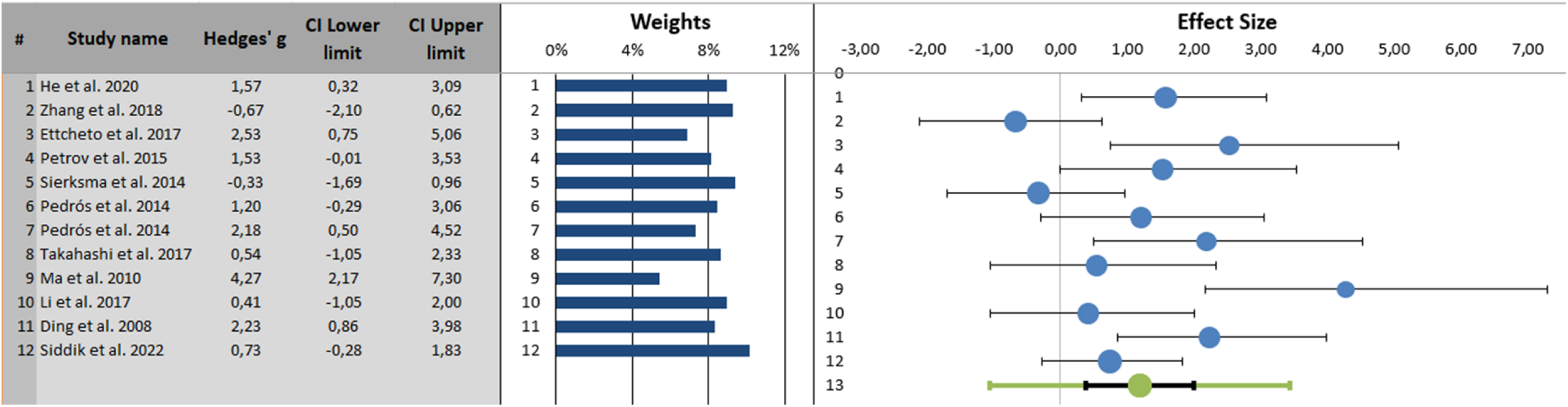
Forest plot of changes in C-terminal domain phosphorylated tau in *APP*_swe_/*PSEN1*_dE9_ vs. control mice. Point estimates of the effect size ± 95% confidence intervals are shown for each study in blue bullets and horizontal lines, respectively. The relative size of each bullet represents a study’s weight in the generation of the meta-analytic result. The green bullet represents the combined effect size (weighted average effect), along with the confidence interval (black horizontal line) and the prediction interval (green horizontal line). The direction of effect size was positive in 10 out of 12 comparisons, indicating significant effects of genotype on the phosphorylation of tau at the C-terminal domain.

**Fig. 12.**
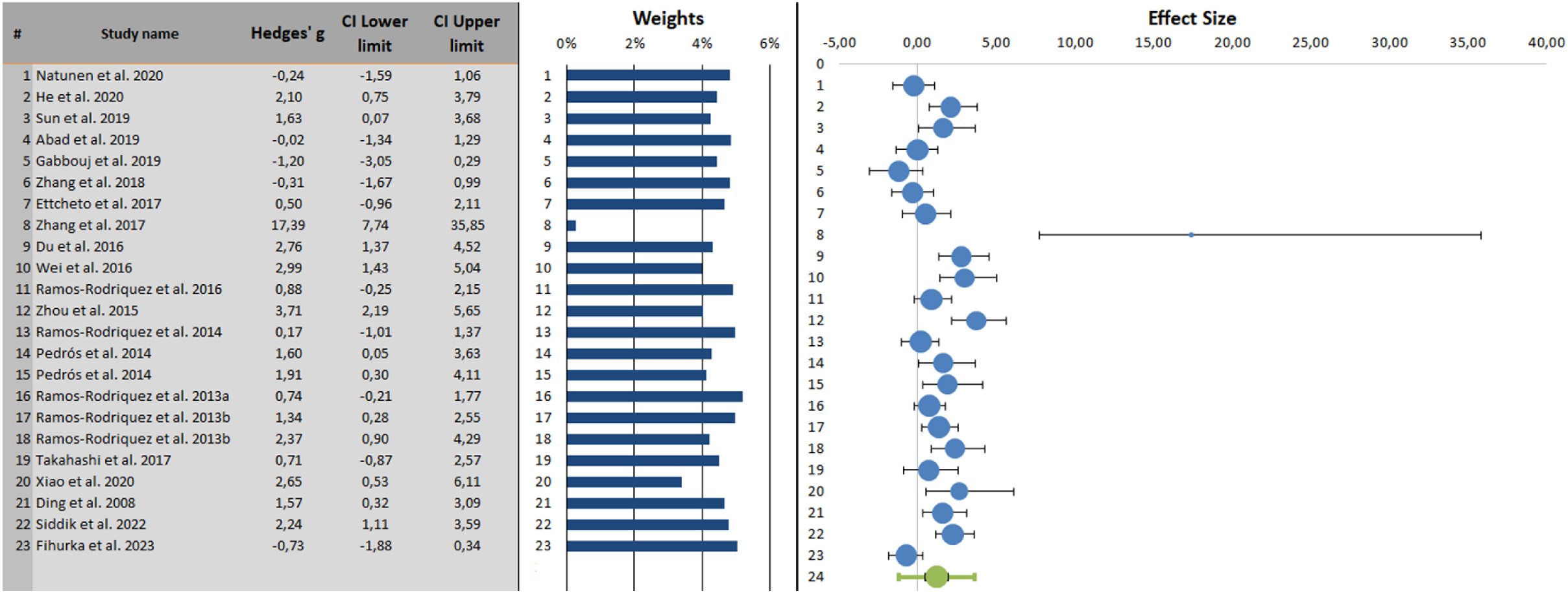
Forest plot of changes in proline-rich phosphorylated tau in *APP*_swe_/*PSEN1*_dE9_ vs. control mice. Point estimates of the effect size ± 95% confidence intervals are shown for each study in blue bullets and horizontal lines, respectively. The relative size of each bullet represents a study’s weight in the generation of the meta-analytic result. The green bullet represents the combined effect size (weighted average effect), along with the confidence interval (black horizontal line) and the prediction interval (green horizontal line). The direction of effect size was positive in 18 out of 23 comparisons, indicating significant effects of genotype on the phosphorylation of tau at the proline-rich domain.

**Fig. 13.**
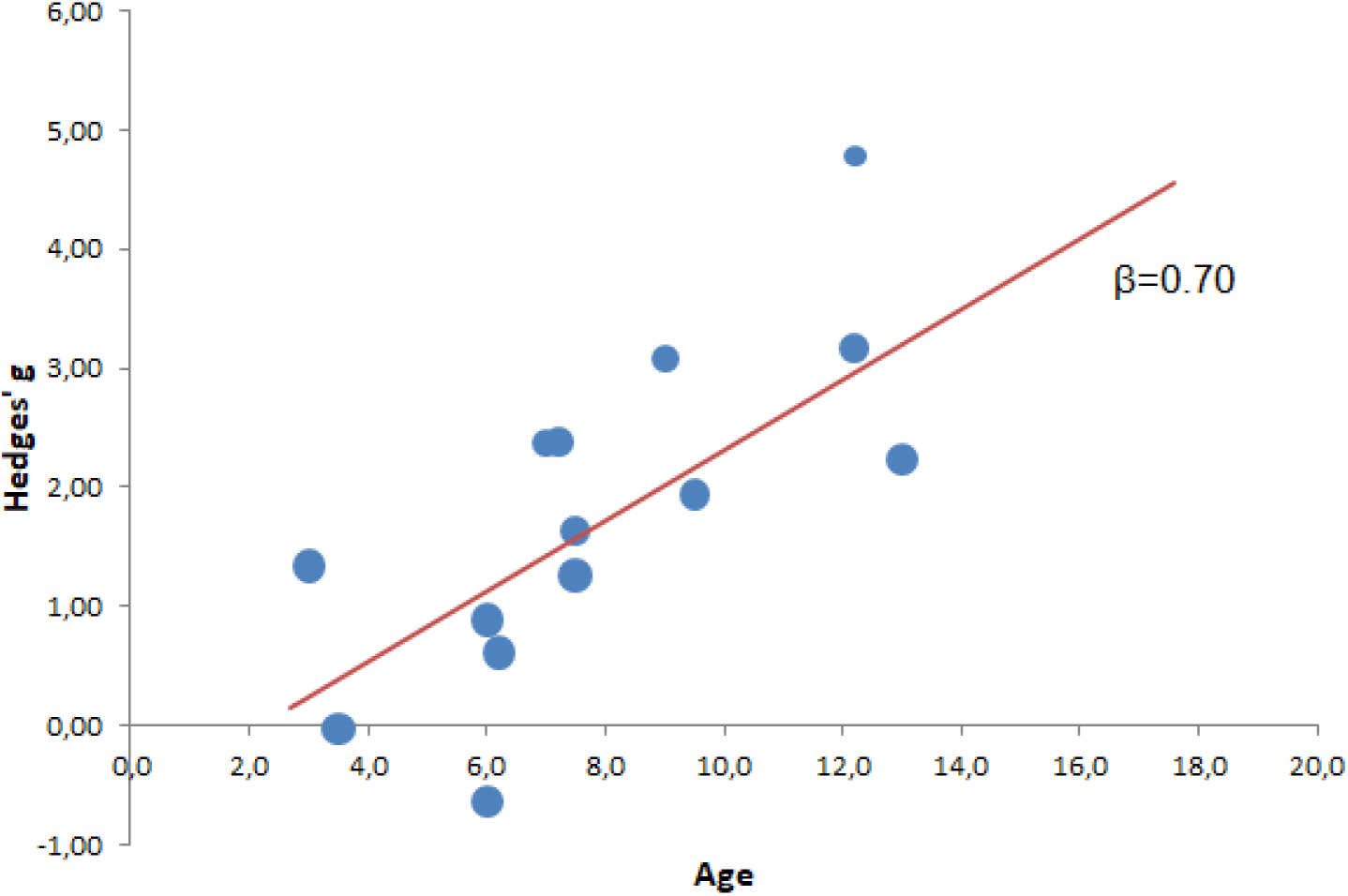
Meta-regression performed with age as moderator in *APP*_swe_/*PSEN1*_dE9_ vs. control mice. Scatter plot and regression line for the moderating effects of age on the phosphorylation of tau at the proline-rich domain in the cortex of *APP*_swe_/*PSEN1*_dE9_ mice. Each data point represents the effect size observed at a specific time-point. There was a progressive increase in the magnitude of hyperphosphorylation in transgenic mice with ageing.

### 3.8 Publication bias: *APP*_swe_/*PSEN1*_dE9_ model

Funnel plot asymmetry was estimated by Egger’s regression. Trim & fill analysis imputed 4, 0 and 1 missing studies for total tau (intercept: 5.94±0.92, t=6.43, *P*<0.000), CT-phosphorylated tau (intercept: 11.35±2.29, t=4.97, *P*=0.001), and PR-phosphorylated tau (intercept: 4.89±1.10, t=4.44, *P*<0.000), respectively. Funnel plots and estimates of adjusted combined effect sizes are shown in Fig. 14A-C.

**Fig. 14.**
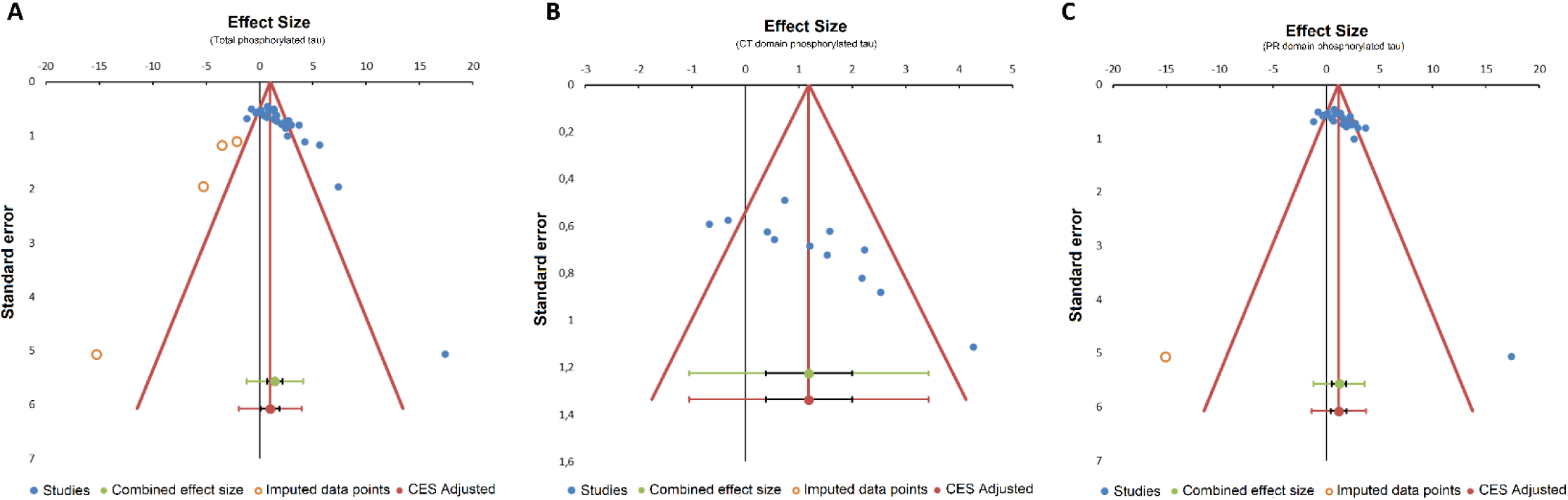
Funnel plot of meta-analysis examining soluble tau phosphorylation in *APP*_swe_/*PSEN1*dE9 vs. control mice. (A) total soluble tau phosphorylation (B) tau phosphorylated at the C-terminal domain (C) tau phosphorylated at the proline-rich domain. Each symbol represents an independent comparison. Adjusted combined effect sizes (CES) were: (A) 1.00±0.42 (CI: 0.15-1.86; PI: -1.96-3.96); (B): 1.18±0.37 (CI: 0.38-1.99; PI: -1.06-3.43); (C): 1.17±0.37 (CI: 0.40-1.94; PI: -1.41-3.75).

### 3.9 Study selection and characteristics: APP23 and J20 mice

For the APP23 model, there were no studies that met the inclusion criteria. For the J20 model, the number of eligible studies was too low to warrant an exploratory meta-analysis (n=6). The flowcharts of study search and selection for APP23 and J20 mice are presented in Supplementary Fig. S6.

## 4. Discussion

The goal of this systematic review was to synthesize and analyse the literature on the phosphorylation of tau protein in mouse models of familial AD. We show that (i) soluble tau is hyperphosphorylated in the brain of 5xFAD and *APP*_swe_/*PSEN1*_dE9_ mice and that (ii) age moderates the association between genotype and tau phosphorylation in a model-specific manner.

Evidence suggests that changes in tau protein phosphorylation, solubility and spreading are pronounced in the brain, not only in AD, but also during physiological ageing, in both humans and experimental animals (Blomberg et al., 2001; Chatterjee et al., 2023; Pettigrew et al., 2022; Sjogren et al., 2001; Torres et al., 2021; Wegmann et al., 2019). In the limited number of studies measuring age-related increases in soluble p-tau in the brain of wild-type mice, levels of phosphorylation were reported to be 1.3- to 3-fold higher in 22 vs. 10-month-old (Bu et al., 2018) and 8 vs. 2-month-old mice (Grinan-Ferre et al., 2016), respectively. These data imply that some degree of tau phosphorylation is inherent to the normal ageing process, and also highlight the pathological nature of the excessive tau phosphorylation that was noted in the meta-analytic comparisons between age-matched transgenic and control animals. Indeed, our analysis revealed consistent and significant effects of genotype on the phosphorylation of soluble tau, supporting the validity of transgenic models in replicating this aspect of tau pathology that is typical of human AD.

Despite the positive direction of effect size among the majority of analyzed studies, there was substantial heterogeneity in the magnitude of tau phosphorylation within both 5xFAD and *APP*_swe_/*PSEN1*_dE9_ mice. This heterogeneity was also apparent when tau pathology was assessed by immunohistochemistry, with some studies, for example, reporting lack of tau-associated immunoreactivity in 5xFAD mice (Oakley et al., 2006; Vergara et al., 2019), and others showing pronounced, age-dependent increases of p-tau in this model (Kang et al., 2021; Neddens et al., 2020). To address the issue of heterogeneity, we applied a random effects model for data analysis, and used subgroup and meta-regression techniques to explore several factors that could be contributing to it, including age, tau domain (proline rich and C-terminal) and brain region (cortex and hippocampus). These factors were chosen *a priori*, to reflect the known spatiotemporal patterns of tau phosphorylation and propagation in the AD brain (Augustinack et al., 2002; Braak and Braak, 1991; Luna-Munoz et al., 2007; Neddens et al., 2018).

In both 5xFAD and *APP*_swe_/*PSEN1*_dE9_ mice, heterogeneity remained high within all studied subgroups, suggesting that factors such as tau domain, brain region, gender, and genetic background, had limited influence on the magnitude of observed p-tau levels. Age, on the other hand, was identified as an important source of heterogeneity in this analysis, since it was shown to moderate the relationship between tau phosphorylation and genotype within certain subgroups. Interestingly, while p-tau levels changed with age in both 5xFAD and *APP*_swe_/*PSEN1*_dE9_ mice, the direction of these changes was different between the two models. In 5xFAD mice, there was a tendency of reduced hyperphosphorylation at the CT domain of tau with advancing age, which reached significance in studies using hybrid mice, female mice, and preparations from the cortex. In the cortex of *APP*_swe_/*PSEN1*_dE9_ mice, however, differences in phosphorylation levels between transgenic and control animals were primarily observed at the PR domain of tau, and tended to increase, rather than decrease, with ageing. These observations suggest that the effects of age on tau phosphorylation are contingent upon the mouse model being examined, emphasizing the importance of tailoring model selection to the appropriate disease stage when assessing the interaction of Aβ and tau. Indeed, considering that all comparisons in our analysis involved age-matched subjects, any differences in the effects of age on the levels of p-tau observed between 5xFAD and *APP*_swe_/*PSEN1*_dE9_ mice are probably linked to variations in the disease stage under investigation, rather than being solely attributed to ageing.

Both 5xFAD and *APP*_swe_/*PSEN1*_dE9_ mice exhibit Aβ deposition in the brain, but with different onset and progression trajectories. Cerebral amyloidosis in 5xFAD mice is aggressive, starting at 2-3 months of age and reaching a plateau in certain areas of the male cortex and hippocampus by as early as 10 months of age (Bhattacharya et al., 2014; Oakley et al., 2006). The onset and progression of amyloidosis in *APP*_swe_/*PSEN1*_dE9_ mice is slower compared to 5xFAD mice, with cortical Aβ accumulation becoming evident by 6 months of age and plateauing by 18 months of age (Babcock et al., 2015; Jankowsky et al., 2004). It is thus, plausible that the progressive reduction in the magnitude of tau hyperphosphorylation at the CT domain, as observed here in 1-13-month-old 5xFAD mice, is driven by the advanced phases of amyloidosis, while the progressive increase in PR phosphorylated tau, as observed in 3-13-month-old *APP*_swe_/*PSEN1*_dE9_ mice, is associated with stages of escalating Aβ pathology. Although speculative, this interpretation is in line with data which show that markers of PR domain phosphorylated tau exhibit close correlations with early Aβ pathology (Mattsson-Carlgren et al., 2020; Ossenkoppele et al., 2021), while CT domain phosphorylated tau is generally considered as a late marker in the AD continuum (Neddens et al., 2018; Stefanoska et al., 2022; Zhou et al., 2006). It is also important to note that longitudinal studies of soluble tau biomarkers in the cerebrospinal fluid (CSF) of familial AD patients, show that the extent of phosphorylation at the PR domain of tau increases significantly in the presence of amyloid pathology, but then decreases as the disease progresses (Barthelemy et al., 2020; Fagan et al., 2014; McDade et al., 2018). Similar patterns of phosphorylation have been observed for the CT domain of soluble tau in cross-sectional studies (Gobom et al., 2022). It is thus reasonable to hypothesize that measuring the levels of p-tau across the entire amyloidosis spectrum in 5xFAD and *APP*_swe_/*PSEN1*_dE9_ mice could reveal a bell-shaped response, in both models.

Although our meta-analysis provides valuable insights into the magnitude of Αβ-induced tau phosphorylation and the effects of age in models of AD, it is essential to acknowledge several potential limitations. First, the exploratory nature and considerable methodological variation that are inherent to preclinical studies, combined with the relatively low number of subjects included in each study, warrant caution in assessing and interpreting the precision of the effect size estimates. This is particularly true for the subgroup analyses, which were performed on an even more limited number of studies, often resulting in imbalanced comparisons between the subgroups. Second, our RoB analysis was inconclusive for the majority of studies, which suggests that there may be sources of heterogeneity not accounted for in our analysis. Third, there is a paucity of studies measuring tau phosphorylation as a primary endpoint in models of familial AD, limiting the available data pool for this meta-analysis. In 5xFAD and *APP*_swe_/*PSEN1*_dE9_ mice, data points on the phosphorylation of tau at the extreme ends of the amyloidosis spectrum are clearly lacking, while the number of studies in J20 and APP23 mice was too low to even attempt to disentangle the effects of *PSEN* mutations on the phosphorylation of tau.

It is thus advisable to approach this study as an exploratory meta-analysis, which is aimed at describing the direction of effect sizes and investigating the causes of heterogeneity, rather than calculating the precise extent of tau phosphorylation in models of AD. That said, it is important to note that Αβ-induced hyperphosphorylation of tau was noted in the majority of analyzed studies, with a combined effect size that remained positive even after adjusting for missing data. Collectively, our observations suggest that the interaction between Aβ and tau in models of AD is dynamic, with the trajectories of tau hyperphosphorylation in transgenic mice corresponding to distinct stages of Aβ-induced pathology. These data imply the presence of optimal intervention points for the administration of both anti-amyloid and anti-tau therapies, highlighting the importance of tailoring model selection to the appropriate disease stage when assessing potential treatments for AD.

## Acknowledgements

The authors would like to extend their gratitude to all contributors who provided raw data from their studies when they were contacted.

## Author contributions

AM was involved in Conceptualization; Methodology; Data curation; Formal analysis; Writing - Original Draft. MK was involved in Methodology; Data curation; Writing - review & editing.

