## Supplementary Files for "A systematic review and meta-analysis of tau phosphorylation in mouse models of familial Alzheimer’s disease"

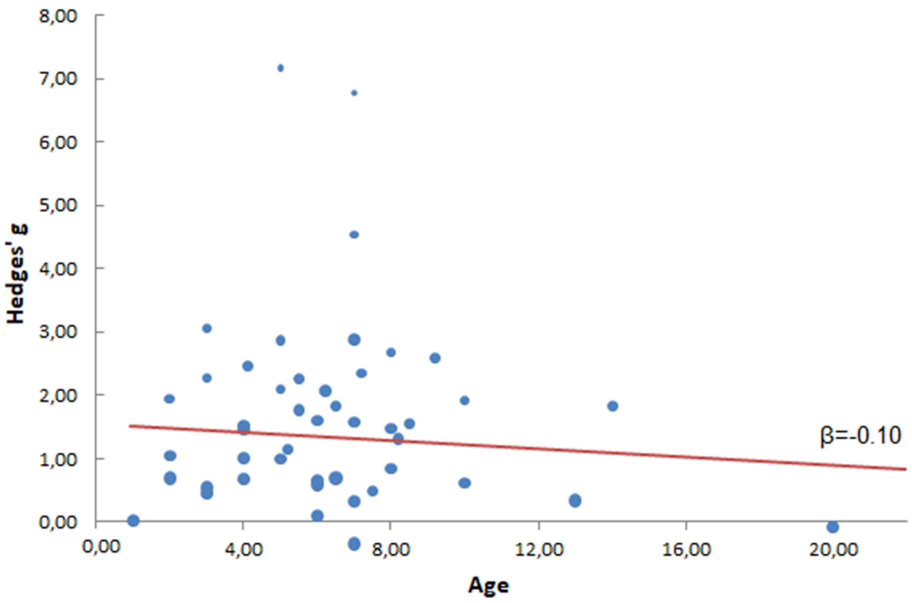


**Supplementary Fig. S1. Moderating effect of age on the levels of soluble p-tau in 5xFAD vs. control mice.**

Scatter plot and regression line for the effects of age on the phosphorylation of total soluble tau across all studies examined. Each data point represents the effect size observed at a specific time-point. Age had no effect as a potential moderator of the relationship between genotype and soluble tau phosphorylation.

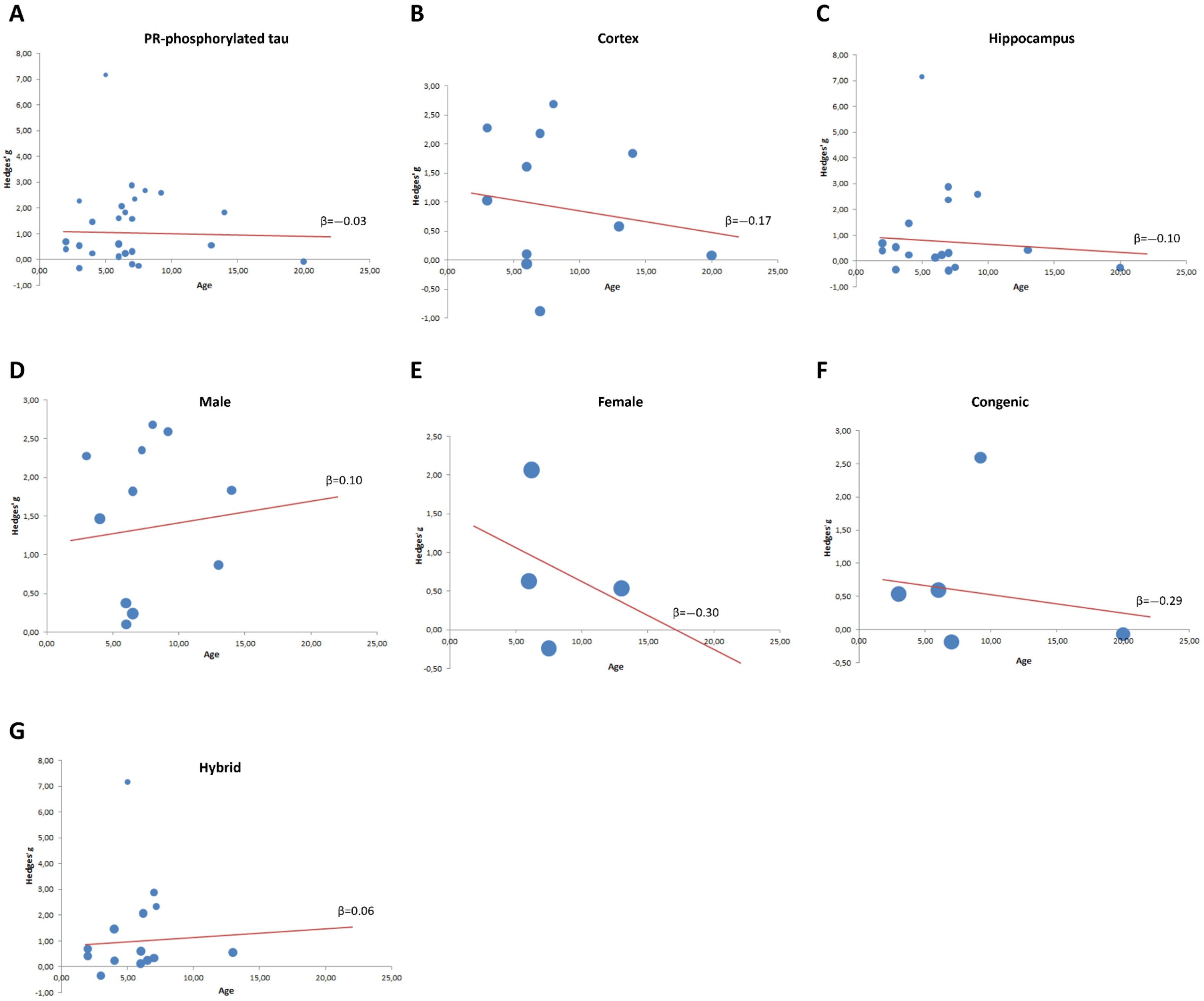


**Supplementary Fig. S2. Moderating effects of age on the phosphorylation of tau at the PR domain in 5xFAD vs. control mice.**

Scatter plots and regression lines for the effects of age on PR domain phosphorylated tau across (A) all retrieved studies (B) studies using preparations from the cortex and (C) the hippocampus (D) studies using male mice (E) female mice (F) mice on a congenic and (G) hybrid background. Each data point represents the effect size observed at a specific time-point. No significant effects of age were observed.

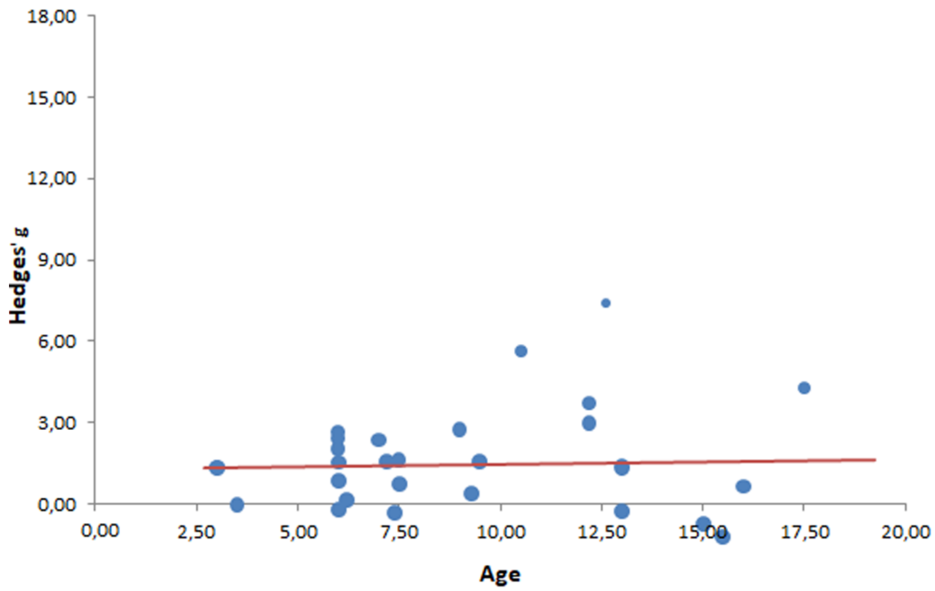


**Supplementary Fig. S3. Moderating effect of age on the levels of soluble p-tau in *APP*$_{swe}$/*PSEN1*$_{dE9}$ vs. control mice.**

Scatter plot and regression line for the effects of age on the phosphorylation of total soluble tau across all studies examined. Each data point represents the effect size observed at a specific time-point. Age had no effect as a potential moderator of the relationship between genotype and soluble tau phosphorylation.

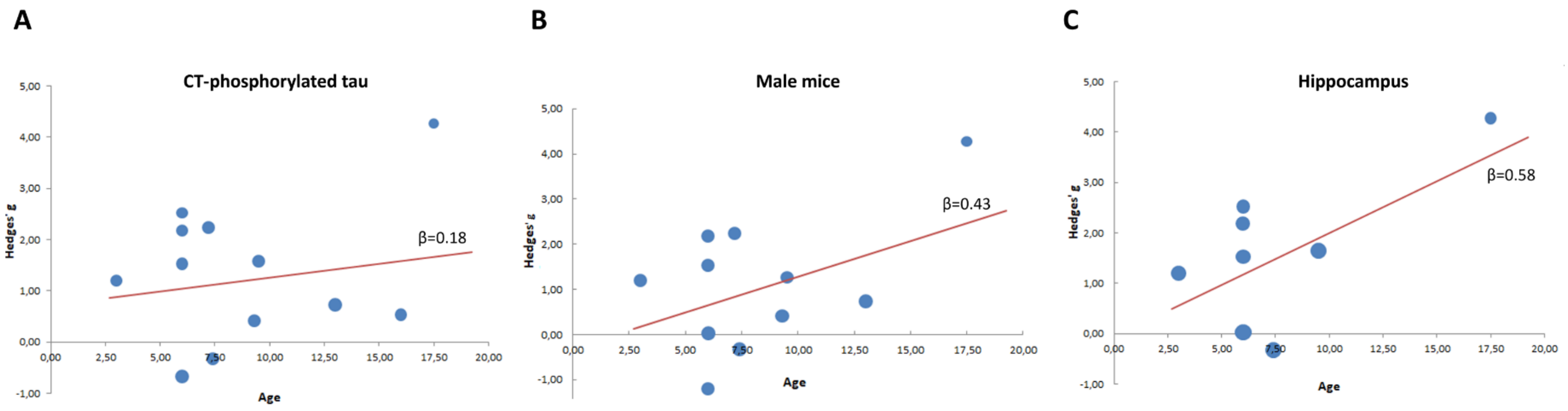


**Supplementary Fig. S4. Moderating effects of age on the phosphorylation of tau at the CT domain in *APP*$_{\text{swe}}$/*PSEN1*$_{\text{dE9}}$ vs. control mice.**

Scatter plots and regression lines for the effects of age on CT domain phosphorylated tau across (A) all retrieved studies (B) studies using male mice and (C) studies using preparations from the hippocampus. Each data point represents the effect size observed at a specific time-point. No significant effects of age were observed.

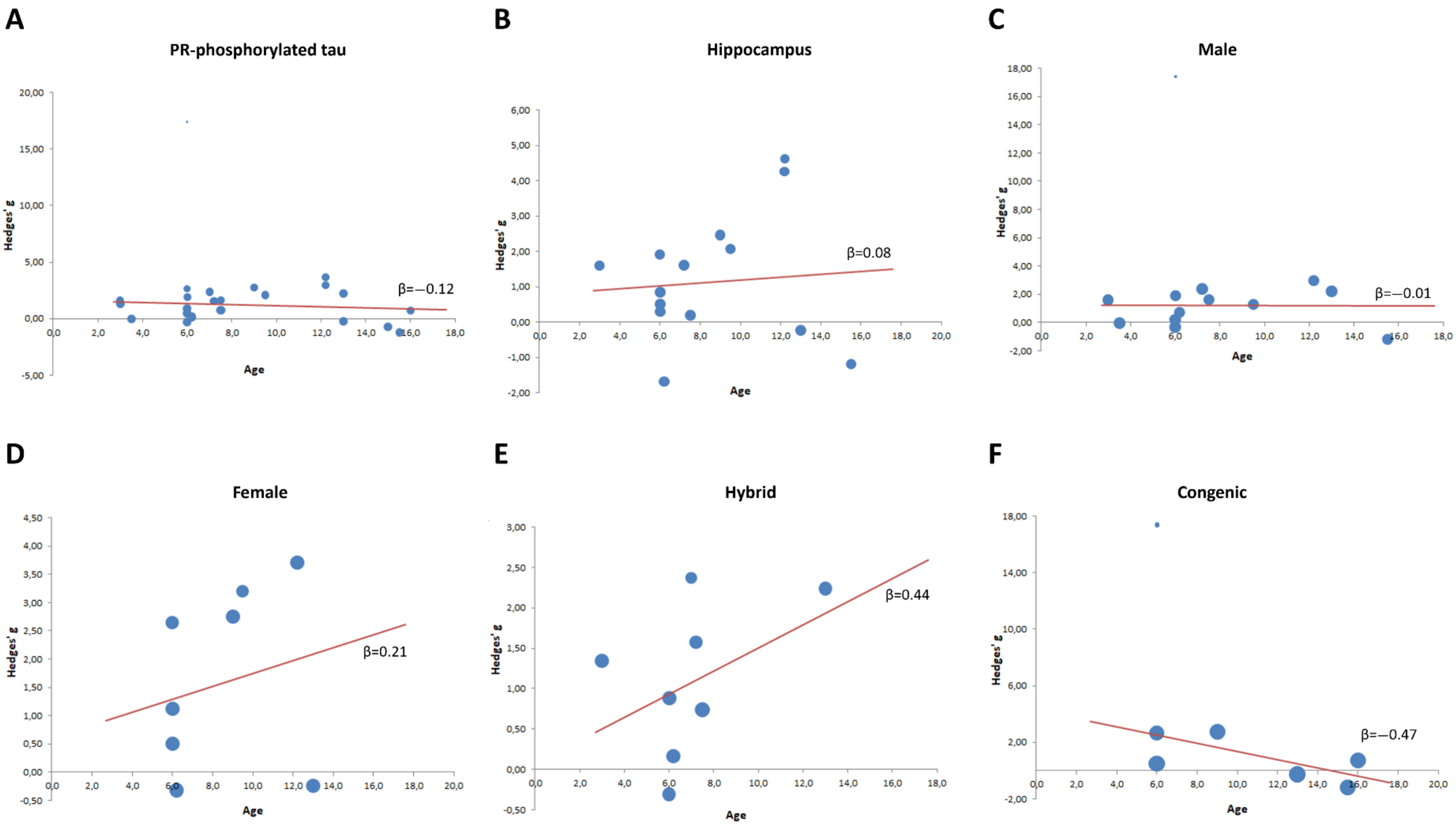


**Supplementary Fig. S5. Moderating effects of age on the phosphorylation of tau at the PR domain in *APP*$_{swe}$/*PSEN1*$_{dE9}$ vs. control mice.**

Scatter plots and regression lines for the effects of age on PR domain phosphorylated tau across (A) all retrieved studies (B) studies using preparations from the hippocampus (C) studies using male and (D) female mice, and studies with mice on (E) a hybrid and (F) a congenic background. Each data point represents the effect size observed at a specific time-point. No significant effects of age were observed.

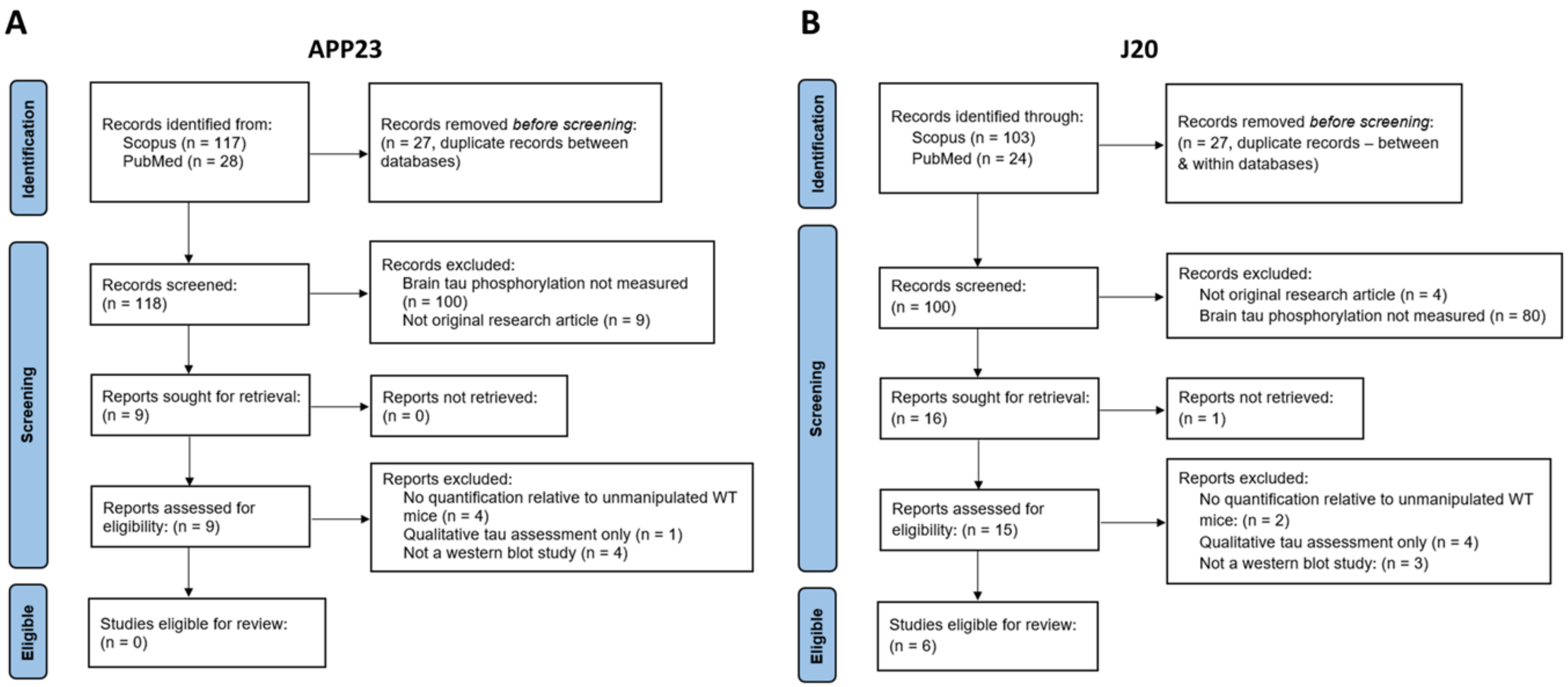


**Supplementary Fig. S6. Flowchart of study search and selection for (A) APP23 and (B) J20 mice.**
